# Nuclear mTOR-Polycomb synergism at developmental gene promoters prone to activation

**DOI:** 10.64898/2026.04.30.721885

**Authors:** Persia Akbari Omgba, Gunwant Patil, Maximilian Stötzel, Chieh-Yu Cheng, Henri Niskanen, Max Ruwolt, Aurora Elhazaz Fernandez, Neil P. Blackledge, Astrid Grimme, Fan Liu, Denes Hnisz, Robert J. Klose, Aydan Bulut-Karslioglu

**Affiliations:** Max Planck Institute for Molecular Genetics, Berlin, Germany; Department of Mathematics and Computer Science, Freie Universität Berlin, Germany; Institute of Chemistry and Biochemistry, Department of Biology, Chemistry and Pharmacy, Freie Universität Berlin, Germany; Leibniz Institute for Molecular Pharmacology, Berlin, Germany; University of Oxford, Department of Biochemistry, South Parks Rd, Oxford OX1 3QU

## Abstract

Pluripotent cells hold the capacity to differentiate into cells with diverse functions, enabling development and regeneration. The balance between local epigenetic repression and maintaining the potential for prompt transcriptional activation supports pluripotency while allowing imminent differentiation. How the activation of developmental pathways is correctly timed in sync with the environment remains poorly understood. Here we find that the cellular growth regulator mTOR selectively binds developmental gene promoters in pluripotent mouse embryonic stem cells. mTOR binding at target genes correlates with and depends on histone H2AK119 monoubiquitination (H2AK119ub1) deposited by the Polycomb Repressor Complex 1 (PRC1). Acute depletion of the whole PRC1 complex or its catalytic activity leads to depletion of mTOR at target gene promoters, whereas forced PRC1 recruitment to an artificial site brings along mTOR. At target genes, mTOR colocalizes with components of the transcription machinery and we find that mTOR-bound genes are distinctly characterized by high levels of RNA Polymerase II pausing. Our findings reveal a role of mTOR in pluripotency regulation and highlight the tight crosstalk between gene regulatory and cell growth machineries in stem cells.

## INTRODUCTION

In vivo, pluripotency is a transient state that only exists for a few days in early mammalian development. Embryonic stem cells (ESCs) derived from blastocyst-stage embryos propagate pluripotency for longer periods in vitro. In the mouse, ESCs grown in the presence of serum and leukemia inhibitory factor (LIF) show transcriptional and epigenetic signatures of the early post-implantation epiblast, while ESCs grown in 2i/LIF capture naïve pluripotency^1,2^.

To ensure acute responsiveness to differentiation signals, developmental genes in ESCs are kept in a repressed, but quickly activatable state. This is achieved by bivalent enrichment of repressive and transcription-associated ‘active’ marks around transcription start sites (TSS)^3–6^. Bivalent TSSs are resolved to monovalent active or repressed states relevant to lineages of choice. Polycomb repressive complexes 1 and 2 (PRC1/PRC2) with their several subcomplexes create partially self-propagating focal heterochromatin environments^7–12^. The catalytic subunit of PRC1, RING1A/B, monoubiquitinates histone H2A lysine 119 (hereafter H2AK119ub1). A subset of PRC2 complexes recognize this mark and deposit trimethyl groups at histone H3 lysine 27 (H3K27me3)^13^. In return, PRC1 complexes recognize H3K27me3 and deposit H2AK119ub1^14^, creating a tight PRC1-PRC2 cross-talk that ensures local gene repression. While developmental Polycomb targets are temporarily repressed, they are protected from permanent repression via antagonism between PRC and DNA methylation complexes^15^. On the other hand, recruitment of RNA Polymerase II (Pol2) to developmental gene promoters and antagonism between PRC1 and Pol2 machineries maintain these genes in a metastable state prone to activation^16,17^. The short half-life of H2AK119ub1 allows prompt clearance of repression and increased Pol2 activity in a lineage-appropriate manner^9,10,16^. Unlike developmental genes, other PRC targets such as metabolic genes^18^ are only lightly repressed, thus presence of neither H3K27me3 nor H2AK119ub1 is a de facto black-and-white indicator of gene silencing.

mTOR is a serine-threonine kinase that is widely known as a central regulator of cell growth that gauges the availability of nutrients and growth factors and adjusts cell growth accordingly^19^. For this, mTOR phosphorylates several target genes that are themselves key regulators of major anabolic processes in the cell such as protein synthesis and energy production. In addition, accumulating evidence suggests that mTOR can function in the nucleus, binding ribosomal DNA and hormone-responsive regulatory elements to promote transcription^20–24^. In both these contexts, the proposed function of nuclear mTOR is related to its cytoplasmic function of growth control.

Along with growth control in numerous physiological and pathological contexts, mTOR activity is essential for the development of early embryos, with *mTOR* KO embryos failing to develop beyond early post-implantation (∼E5.5)^25^. In comparison, ribosomal growth disorders are not necessarily embryonic lethal and when they are, the embryos fail at mid-to late-gestation^26,27^. Therefore, the early lethality of mTOR KO embryos imply functions that go beyond growth and metabolic regulation. Furthermore, inhibition of mTOR activity (**mTORi**) in blastocyst-stage embryos temporarily and reversibly pauses further development, placing mTOR as a regulator of developmental timing^28,29^. mTORi-treated mouse blastocysts and human blastoids maintain naïve/formative pluripotency for longer periods and show the capacity to further develop once inhibition is lifted^29–31^. Thus, under mTORi, pluripotency is stabilized in vivo for longer than otherwise possible.

Here we identify a role for mTOR as a chromatin binder in pluripotent mouse ESCs. Acute perturbation assays reveal synergism between mTOR and PRC1 activity and place mTOR in the PRC-Pol2 axis at developmental genes.

## RESULTS

### mTOR perturbations affect the pluripotent state of mouse ESCs

Analogous to the stabilization of pluripotency in vivo during mTORi-induced dormancy, we previously observed that mTORi further stabilizes pluripotency of mouse ESCs cultured in serum/LIF condition, yielding ESC colonies that are dormant but uniformly highly express alkaline phosphatase (AP) and the stem cell surface antigen SSEA1^32^. To further probe the link between pluripotency and mTOR activity, we generated a mouse ESC line that allows genetic deletion of the entire mTOR locus in an inducible manner (hereafter **mTOR iKO**, Figure S1a-c). mTOR iKO ESCs carry loxP sites 5’ and 3’ to the mTOR gene, which are recombined in the presence of a Cre recombinase (Figure S1a, b). For the deletion, we nucleofected Cre-GFP and sorted cells after 48h. Deletion was confirmed by genotyping (Figure S1b). We observed, however, that mTOR KO cells were depleted from culture within the next 2-3 days, with protein expression and cellular proliferation increasing again after a split (Figure S1d). This is likely due to the lethality of mTOR homozygous deletion. No homozygous KO colonies were recovered after sorting single cells after nucleofection (data not shown). Alternatively, we inserted tamoxifen-translocatable CreERT2 transgene in mTOR^loxP/loxP^ cells into the *H11* locus. Despite successful integration into different safe harbor loci, monoclonal selection, and expression of CreERT2, mTOR could not be uniformly depleted (data not shown). Finally, as an alternative approach, we knocked-in a degron tag (dTAG) at the mTOR locus both homozygously and heterozygously (Figure S1e, f). While heterozygous degradation reliably worked, homozygous degradation was not successful (Figure S1g). The former has no effect on ESC fitness, morphology or proliferation, which aligns with embryonic mTOR hypomorphs not altering fitness or development^25^. We concluded that complete depletion of the mTOR protein is incompatible with ESC culture and that the mTOR locus is heavily regulated to counteract depletion.

To nevertheless address a potential role of mTOR in pluripotency regulation, we opted to collect and analyze mTOR iKO cells within 2 days of nucleofection. Like mTORi-treated ESCs, mTOR iKO ESCs reduce proliferation, form uniformly round colonies and increase SSEA1 expression (Figure 1a-b, S1d). Thus, suppression of mTOR reinforces pluripotency in mouse ESCs. Unlike mTORi however, mTOR iKO ESCs do not form a stable culture and cannot be propagated, suggesting that minimal mTOR activity is required to maintain viability.

**Figure 1.**
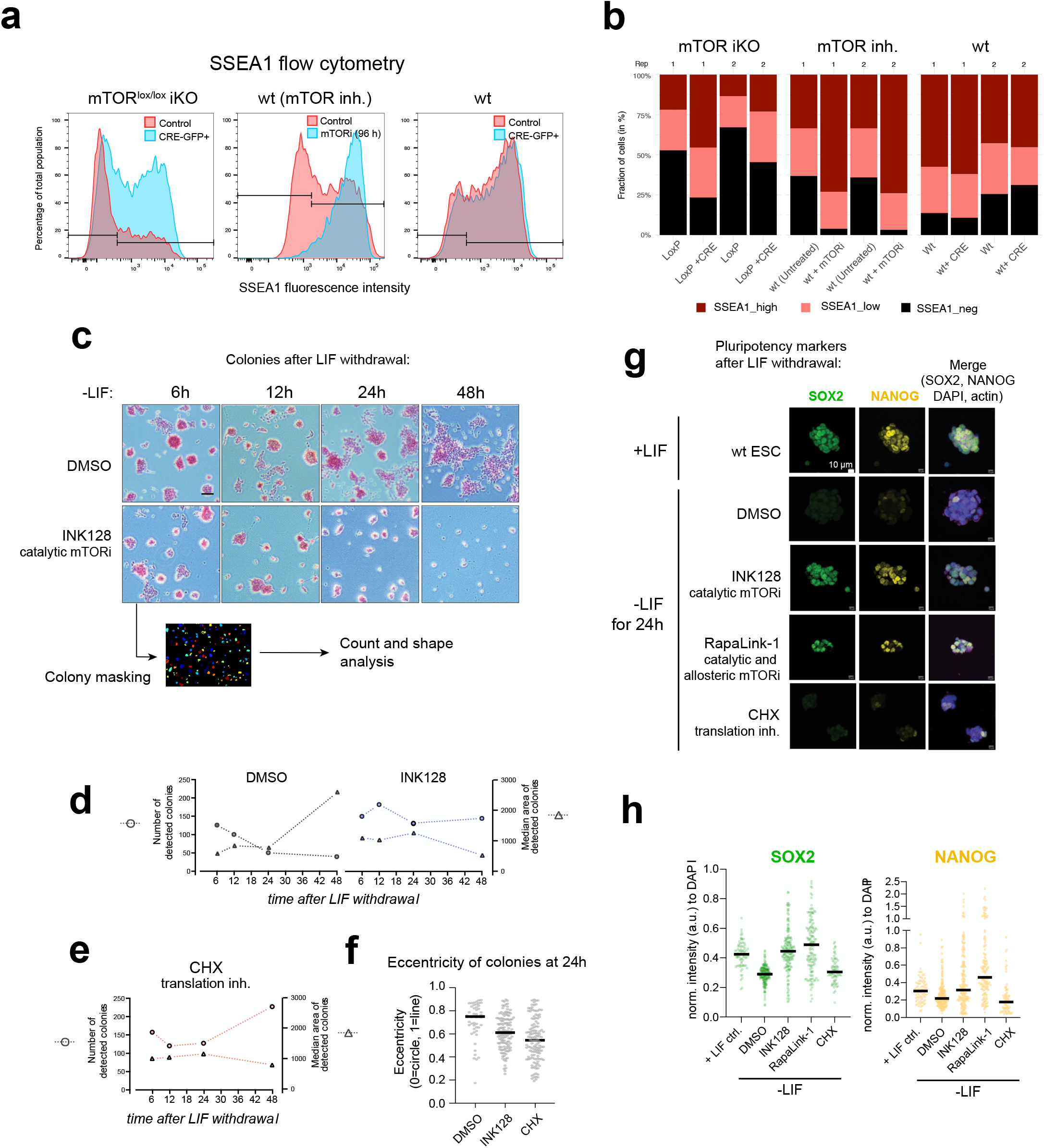
mTOR perturbations affect the pluripotent state of mouse ESCs. a. Flow cytometry analysis of the stem cell surface marker SSEA1 in mTOR iKO, mTORi, and wild-type conditions. b. Quantification of SSEA1-negative, low, and high populations in biological replicates. c. Alkaline phosphatase staining of mouse ESCs 6-24h after LIF withdrawal with or without mTORi (INK128) treatment. Images were analysed using Cell Profiler to count colonies and measure morphological parameters. d. Number (left) and area (right) of detected colonies (left) in DMSO and mTORi conditions. e. Number (left) and area (right) of detected colonies (left) in CHX (translation inhibition) condition. f. Eccentricity (circularity) of detected colonies in the indicated conditions. g. Representative images of immunofluorescence staining of the pluripotency markers SOX2 and NANOG in ESCs (in IF) vs 24h after LIF withdrawal. Cells were treated with two different mTORi or CHX. DMSO serves as control. h. Quantification of single cell SOX2 and NANOG fluorescence intensities. Each dot corresponds to a cell and horizontal line denotes median intensity.

Over-reinforcement of the pluripotency network often compromises the ability to timely exit from pluripotency and the responsiveness of ESCs to differentiation cues^33,34^. Therefore, we next tested whether mTOR-perturbed cells are resistant to exit from pluripotency. For this, we cultured ESCs without leukemia inhibitory factor (LIF) for up to 48h with or without the mTORi INK128 (Figure 1c). Withdrawal of LIF allows wild-type ESCs to promptly exit from pluripotency, evidenced by disrupted and enlarged colonies and loss of SOX2 and NANOG expression (Figure 1c-h). In contrast, mTORi-treated cells maintained colony morphology and SOX2 and NANOG expression 24h after LIF removal. This effect is likely not due to reduced protein synthesis capacity, as ESCs treated with the translation inhibitor cycloheximide (CHX) could exit pluripotency in this time frame (Figure 1h) despite showing morphological similarities to mTORi colonies (Figure 1e, f). We thus conclude that suppression of mTOR activity poses a barrier against pluripotency exit.

We next tested whether ectopically high mTOR expression may tip the balance against pluripotency and force ESCs to differentiate. For this, we overexpressed either wild-type, hyperactive, or kinase-dead full length mTOR in ESCs via a piggybac expression system (Figure S2a)^35^. Cells were sorted for mCherry twice in a 6-day period (to counteract silencing of the transgene), cultured in the presence of LIF, and assayed for alkaline phosphatase activity and colony morphology (Figure S2b-e). Overexpression of mTOR in ESCs led to differentiation of colonies, while the kinase-dead version largely rescued this effect (Figure S2c). These results reveal that pluripotency networks are sensitive to increased mTOR expression even though hypomorphs do not adversely affect stemness. Taken together, these mTOR loss- and gain-of-function perturbations suggest that mTOR activity influences the pluripotent state in addition to adjusting cell growth and proliferation levels.

### Nuclear mTOR binds developmental genes in a stage-specific manner

Due to the tight crosstalk with mTOR activity and ESC state, and since mTOR is found in diverse cellular fractions including the nucleus in ESCs (Figure S3a), we hypothesized that mTOR may serve a role in the nuclear regulation of pluripotent cells. To test whether mTOR binds chromatin, we performed chromatin immunoprecipitation followed by sequencing (ChIP-seq) in ESCs and mouse embryonic fibroblasts (MEFs) (Figure 2a-b). We detected considerable mTOR binding at chromatin in ESCs but not MEFs (438 peaks in serum/LIF ESCs vs. 35 peaks in MEFs). In ESCs, mTOR binding at chromatin is restricted to the formative state enacted by serum/LIF culture condition and was minimal in the naïve state in 2i/LIF medium (55 peaks, Figure 2a-b). Target sites in serum/LIF ESCs were identified as those detected robustly across all 5 replicates and mostly comprise of promoters (∼75%, Figure 2c, d, S3b-d). Notably, mTOR-target genes are highly enriched for early developmental regulators and include transcription factors (TFs) across Hox, Pax, Sox, Fox families (Figure 2e, f).

**Figure 2.**
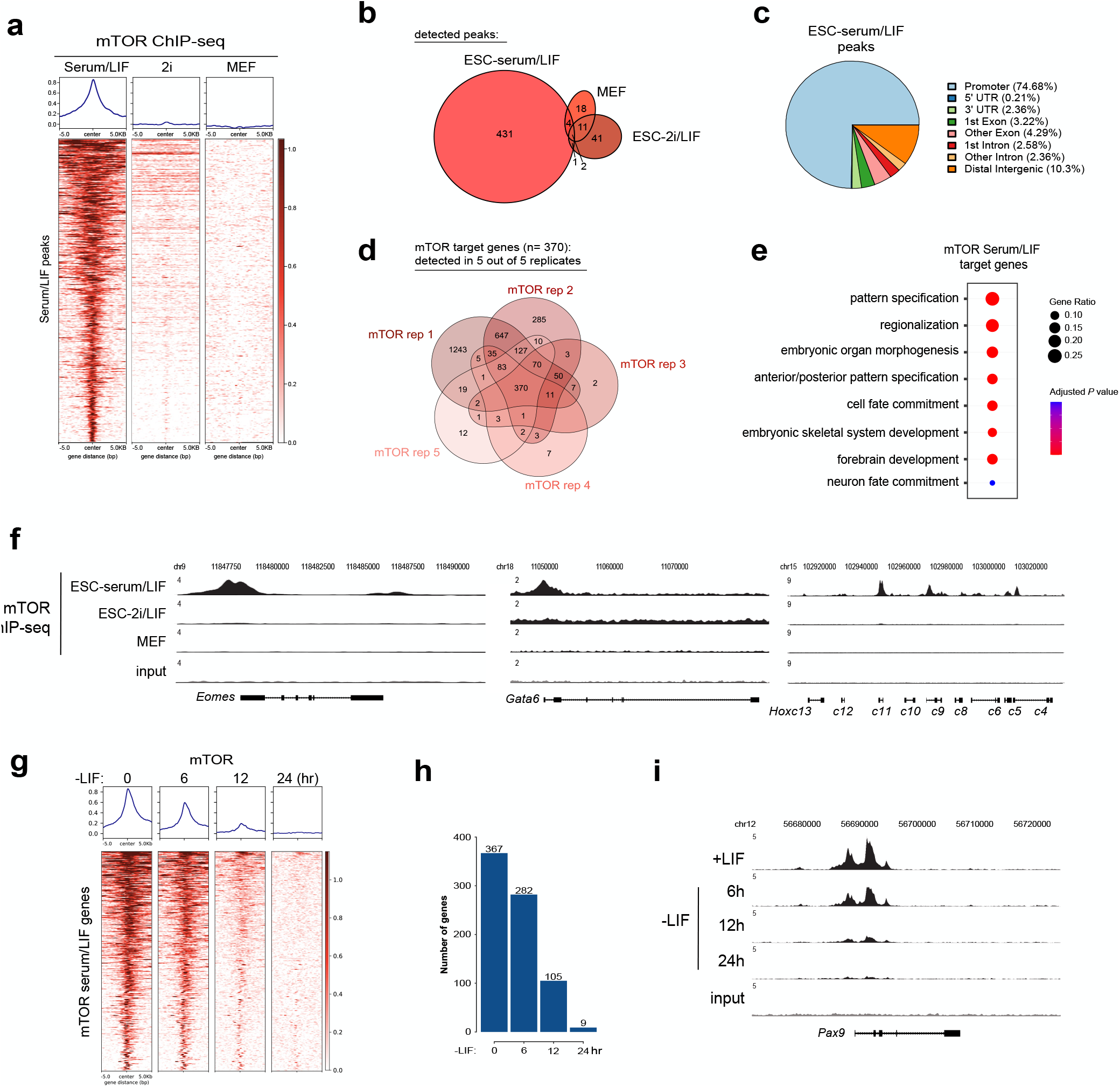
Nuclear mTOR binds developmental gene promoters in serum/LIF ESCs. a. Heatmap showing mTOR occupancy +/-5 kb of its serum/LIF target peaks as identified by MACS2 analysis of ChIP-seq data. b. Venn diagram showing the overlap of peaks in ESC-serum/LIF, ESC-2i/LIF, and mouse embryonic fibroblasts (MEF). c. Pie chart showing genomic distribution of mTOR serum/LIF peaks. d. Venn diagram showing the identification of mTOR target genes. Only genes detected in all 5 replicates were considered true targets. For QC plots, see Figure S3. e. Gene ontology analysis of mTOR peaks in serum/LIF ESCs. Only promoter peaks were used. f. Genome browser snapshots of mTOR binding profiles in shown conditions. g. Heatmap showing mTOR ChIP-seq signal +/-5 kb of the TSS of target genes within 24h of LIF withdrawal. h. Number of mTOR target genes identified in samples shown in (G). i. Genome browser snapshot of an mTOR target gene during the first 24h of LIF withdrawal.

Given that mTOR is detected at chromatin in ESCs and not in MEFs, we next tested whether acute differentiation of ESCs leads to depletion of mTOR from chromatin. For this, serum/LIF ESCs were allowed to exit pluripotency via withdrawal of the pluripotency safeguarding factor LIF. mTOR remained bound at target genes until after 6h of LIF withdrawal and became gradually depleted from most targets between 6-24h (Figure 2g-i). Together, these results show that mTOR binds gene promoters – directly or indirectly – in serum/LIF ESCs until after exit from pluripotency.

### Nuclear mTOR targets two sets of genes with distinct epigenetic makeup

We next surveyed the chromatin features of mTOR target genes to find out potential recruitment mechanisms. A broad gene ontology analysis assigned mTOR target genes to developmental pathways (Figure 2e). Developmental genes are temporarily repressed in pluripotent cells through the co-enrichment (i.e. bivalency) of repressive (H3K27me3, H2AK119ub1) and transcription-associated (H3K4me3) histone modifications. We thus profiled the canonical bivalency marks at mTOR target genes (Figure 3a). mTOR targets show a heterogeneous pattern, in which regions that show high levels of mTOR binding correlate with H3K27me3 and H2AK119ub1, and those that show low mTOR levels correlate rather with H3K4me3 (Figure 3a). At individual loci, mTOR signal is correlated most with H2AK119ub1 and anti-correlated with H3K4me3 (Figure 3b). H3K27me3 and H2AK119ub1 coexist at many target genes and show the overall highest correlation in this analysis as expected (Figure 3b). At target gene promoters, mTOR is more focally distributed around the transcription start site (TSS) with some enrichment in the gene body, compared to H2AK119ub1 and H3K27me3, which cover large domains (Figure 3c). A H2AK119ub1-sensitive gene ontology analysis partitions mTOR target genes into two clusters, in which H2AK119^high^ mTOR targets (n=280) associate with developmental pathways and H2AK119^low^ mTOR targets (n=90) associate with metabolic processes (Figure 3d). Based on this analysis and the fact that mTOR binding shows the highest correlation with H2AK119ub1, we sub-clustered mTOR target genes into two clusters according to their H2AK119ub1 levels (Figure 3e-f) and focus on cluster 1 in the remainder of the study.

**Figure 3.**
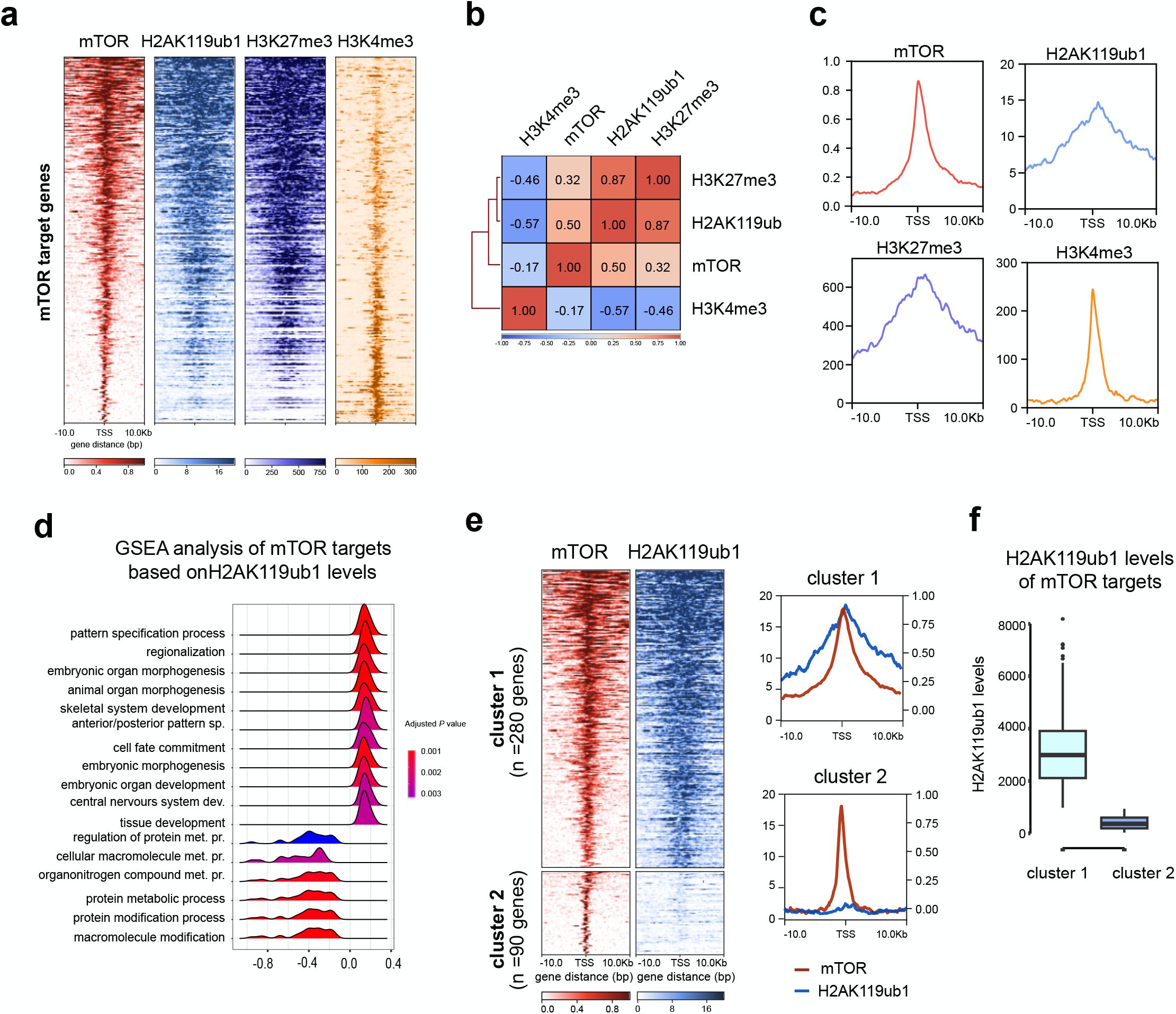
Nuclear mTOR binding is correlated with PRC1-deposited H2AK119ub1. a. Heatmap showing levels of indicated histone modifications at mTOR-target genes. b. Pearson correlation between mTOR and histone modification signals. mTOR binding is most highly correlated with H2AK119ub1. c. Occupancy pattern of mTOR and shown modifications around target gene TSSs. mTOR is focally bound around the TSS. d. Gene ontology enrichment analysis of mTOR target genes, in relation with their H2AK119ub1 levels. e. Left, hierarchical clustering of mTOR-target genes based on H2AK119ub1 levels. Right, distribution of binding signals around TSSs in each cluster. f. Quantification of mTOR and H2AK119ub1 levels in clusters 1 and 2. Lines denote median, the upper and lower edges of the box show the first and third quartiles.

### Nuclear mTOR recruitment at target developmental genes is dependent on H2AK119ub1

Having observed the correlation between mTOR and Polycomb-deposited H2AK119ub1 and H3K27me3 marks, we next tested whether mTOR binding is dependent on the presence of these at chromatin. H2AK119ub1 and H3K27me3 marks are deposited by Polycomb repressive complexes 1 and 2 (PRC1 and 2), respectively. To test causality, we used degron-mediated perturbation to selectively and quickly deplete the PRC1 or PRC2 complex (Figure 4a). For PRC1 degradation, we homozygously inserted degron tags into the endogenous *Ring1a* and *Ring1b* genes via Cas9-assisted gene editing to yield a *Ring1a/1b*^dTAG^ ESC line (Figure S4a). Treatment of this line with the dTAG13 ligand for 3h results in the degradation of the RING1A/1B ubiquitin ligases by the proteasome and depletion of H2AK119ub1 (Figure 4b). For PRC2 depletion, a previously published *Suz12*^*dTAG*^ ESC line^36^ was used to degrade the PRC2 complex upon 3h dTAG treatment. Acute loss of PRC1, but not PRC2, quickly depleted mTOR from chromatin at cluster 1, but not cluster 2, target genes (Figure 4c-d). For comparison, we curated a control gene set (Figure S4b-d) that includes gene promoters that are bound by PRC1, but not mTOR, and carry high levels of H2AK119ub1 (see Methods). mTOR binding at these sites, as expected, were not affected by dTAG treatment (Figure 4c).

**Figure 4.**
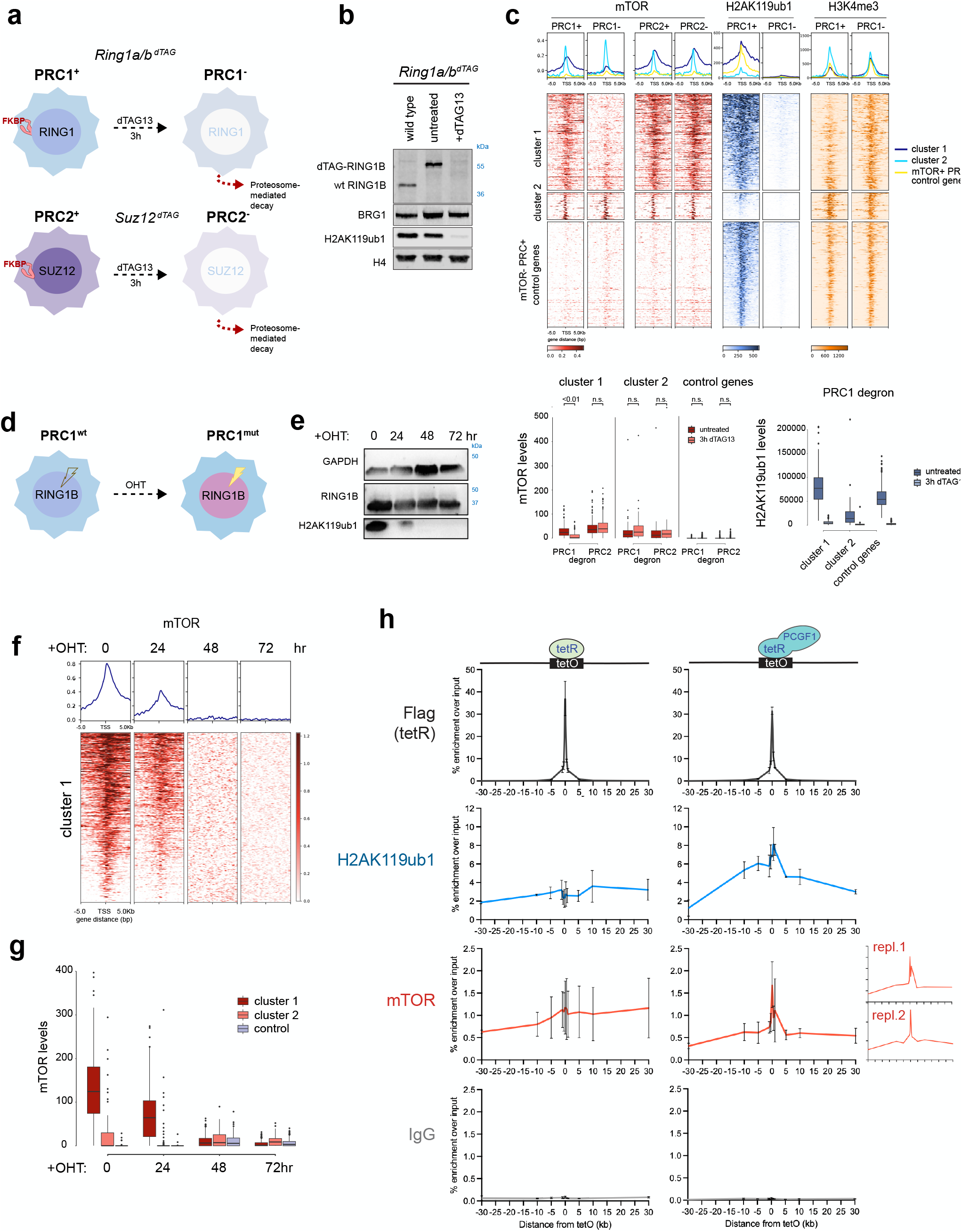
Nuclear mTOR binding at cluster 1 targets is dependent on H2AK119ub1. a. Schematic of PRC1 and PRC2 degron approaches used in the study. For details of genetic constructs, see Figure S4A. b. Western blot showing depletion of RING1B and H2AK119ub1 after 3h dTAG treatment. c. Heatmap showing binding levels +/-5 kb of the TSS of mTOR-target genes and control genes in PRC1 and PRC2 degron cell lines. Control genes sampled from mTOR-negative PRC-bound genes (for details see Figure S4). mTOR recruitment is selectively lost in cluster 1 in PRC1-but not PRC2-ESCs. Bottom, quantificatitative analyses. Lines denote median, the upper and lower edges of the box show the first and third quartiles. d. Schematic of PRC1 inducible catalytic mutant approach used in this study (previously published in Blackledge et al^10^). For details of genetic constructs, see Figure S4. e. Western blot showing gradual depletion of H2AK119ub1 upon OHT treatment. f. Heatmap showing mTOR binding levels +/-5 kb of the TSS of target genes during 72h of OHT treatment. For control genes, see Figure S4f. g. Quantification of mTOR binding signal at mTOR target gene clusters and control genes. h. Deposition of H2AK119ub at an artificial operon results in recruitment of nuclear mTOR. Left, negative control experiment where tetR is expressed without any fusion. Right, the fused tetR-Pcgf1 gene is expressed and is recruited to the tetO site. Plots show % enrichment over input from two biological replicate experiments.

Having observed the dynamic dependency of mTOR on PRC1, we next aimed to pinpoint whether mTOR binding depends on the presence of the PRC1 complex or the H2AK119ub1 mark that it deposits. For this, we used a previously published inducible PRC1 catalytic mutant system, in which, tamoxifen (OHT) treatment leads to replacement of a wild-type exon of *Ring1b* with a catalytic mutant exon via inversion^10^ (Figure 4d and S4e). This catalytically inactive RING1B protein integrates into the PRC1 complex and binds usual target sites^10^. The catalytic mutation of RING1B results in gradual depletion of H2AK119ub1 from chromatin over time without depleting RING1B until at least 72h (Figure 4e). Following this catalytic mutation, mTOR binding at developmental gene promoters mirrored H2AK119ub1 dynamics, whereby both were depleted at cluster 1 targets by 48h (Figure 4f-g, S4f). These results reveal acute dependency of mTOR recruitment on H2AK119ub1 at cluster 1 target genes.

To further test whether H2AK119ub1 is sufficient for recruitment of nuclear mTOR, we employed artificial recruitment of PRC1 to a synthetic bacterial tet operator array (tetO, <1 kb) flanked on both sides by bacterial artificial chromosome sequences and inserted into chromosome 8 in mouse ESCs^43^. The tetO array sequence lacks CpGs and is devoid of gene regulatory elements, therefore has no resemblance to natural targets of the Polycomb complex or mTOR, allowing us to test sufficiency. In this system, we used a tetR-Pcgf1 fusion construct, which was previously shown to effectively deposit H2AK119ub1 at the tetO array^43^. We confirmed recruitment of both tetR-PCGF1 and tetR alone to the tetO array via ChIP-qPCR (Figure 4h). In this setup, tetR-PCGF1 recruitment results in H2AK119ub1 deposition at the tetO array, which then appears to spread to flanking DNA. We could detect mTOR binding in tetR-PCGF1-expressing cells at and around the tetO array at an expected lower level compared to H2AK119ub1 (Figure 4h). mTOR did not spread to the flanking regions unlike the histone mark itself. These results point to nuclear mTOR-PRC1 synergism also outside the natural context of gene regulatory sequences and suggest sufficiency. However, since we did not observe robust mTOR binding at all PRC1 target promoters (Figure S4b), we conclude that H2AK119ub1 is necessary but not sufficient for mTOR recruitment in the natural context and/or mTOR may be removed from some target sites in competition with other binders.

### mTOR interacts with RNA Polymerase II at target genes

To understand the principles behind selective mTOR binding/retention at a subset of PRC1 target gene promoters, we set out to uncover features that distinguish mTOR+ and mTOR-Polycomb target genes. By surveying H2AK119ub1 abundance across all PRC1 targets, we found that mTOR binds those with the highest H2AK119ub1 levels (Figure S4c) and lowest base expression level in ESCs (Figure 5a). It was previously shown that PRC targets are not solely comprised of silenced developmental targets but include genes with other, e.g. metabolic, functions^37^. mTOR binding evidently distinguishes the more repressed developmental targets from the more transcriptionally-pervasive ones. Despite the heavier repression, mTOR+ PRC+ targets are promptly induced when PRC1 is degraded, suggesting that they are poised for activation and that mTOR may contribute to the timing of their activity (Figure 5b, data from Dobrinic et al^16^).

**Figure 5.**
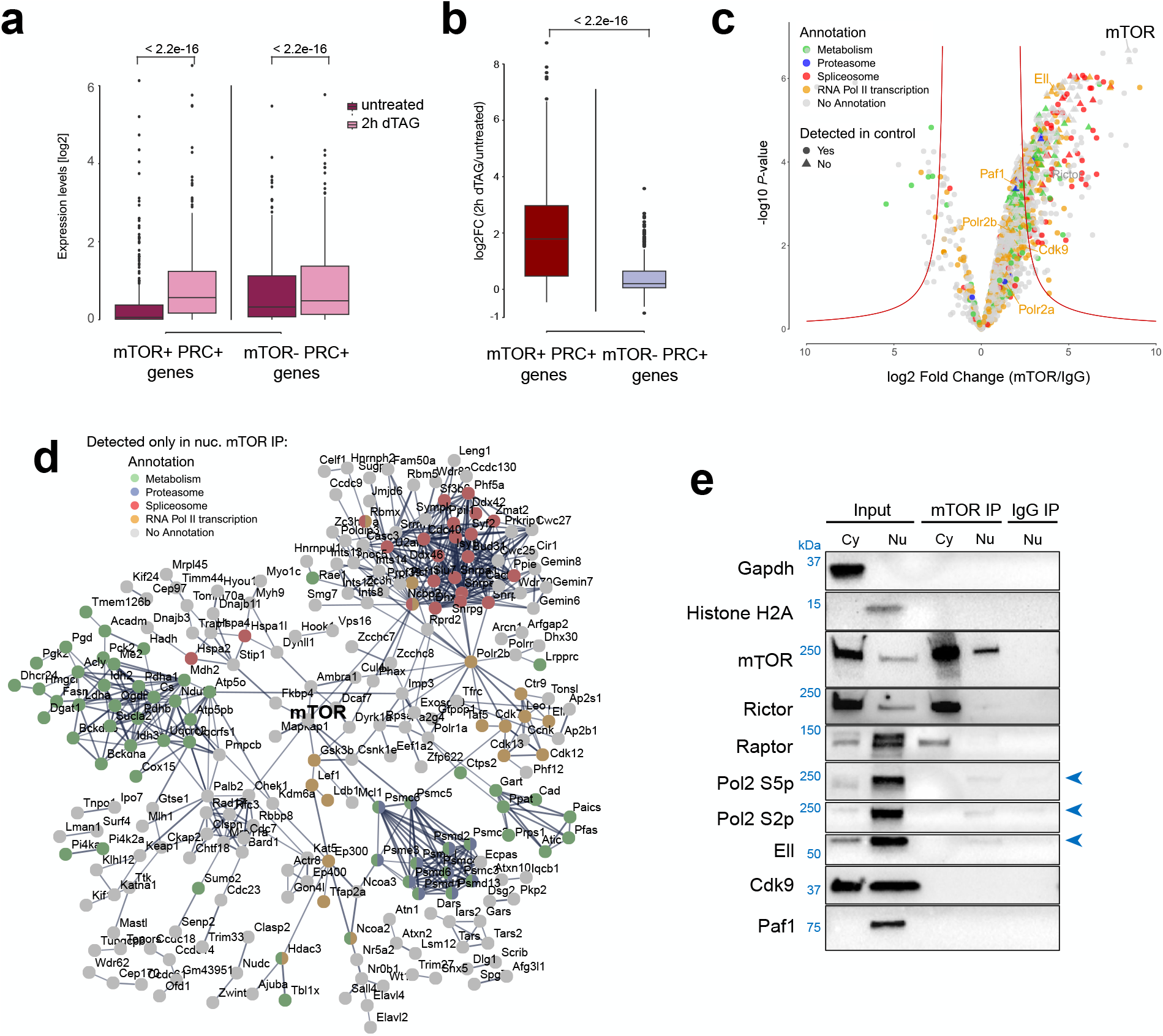
Nuclear mTOR associates with RNA Pol2 complexes. a. Expression level of mTOR+ and mTOR-PRC targets in untreated and PRC1-degraded ESCs. P values were computed by performing paired Wilcoxon Ran Sum tests. b. Log2FC of the expression levels of mTOR+ and mTOR-PRC targets in PRC1-degraded ESCs relative to untreated ESCs. The p values was computed by performing an unpaired Wilcoxon Ran Sum test. c. Volcano plot of proteins detected in nuclear mTOR IP-mass spectrometry. ESCs were fractionated into nuclear and cytoplasmic fractions and the nuclear fraction was used for the IP. Nuclear IgG IP was performed as control. Proteins only detected in mTOR IP are denoted as triangles. Proteins in enriched pathways are colored. d. STRING protein-protein interaction analysis of proteins only detected in mTOR IP (and not in IgG). Pathway enrichment analysis was performed to detect enriched interactions. e. IP-WB validation of mass spectrometry results. Both nuclear and cytoplasmic fractions were used for mTOR IP for comparison reasons.

As a canonically cytoplasmic protein, knowledge about mTOR nuclear interactors in the context of early development is lacking. To address this and identify potential nuclear interactors in ESCs, we next performed nuclear mTOR IP-mass spectrometry. Serum/LIF ESCs were first fractionated into cytoplasmic and nuclear compartments and the nuclear lysate was used for mTOR IP, followed by mass spectrometry (Figure 5c, 4 biological replicates were performed). mTOR itself was identified among the most significant hits and was only detected in mTOR IP and not in IgG, validating our approach (Figure 5c). Protein-protein interaction analysis revealed and pathway enrichment analysis revealed that proteins with metabolic, splicing, proteosome, and transcription regulatory functions are among those that co-precipitated with mTOR (Figure 5d, S5a, b, Figure S5 shows all proteins over log2FC>2 and p value<0.05 and Figure 5d shows those only detected in mTOR IP and not in IgG). Spliceosome proteins were among the highest enriched proteins in mTOR IP over IgG IP, however were often also detected in IgG IP and were flagged as potentially spurious hits (Figure 5c, S5b).

Among potential nuclear mTOR interactors, we focused on transcriptional regulators due to the prompt activatability of mTOR-target genes. Among these, RNA Pol2 components (Polr2a, 2b), Cdks 9, 12, and 13, Paf complex proteins Paf1 and Leo1, and the elongation regulator Ell were categorized as of highest relevance. To validate whether nuclear mTOR indeed interacts with these proteins, we performed IP-WB on cytoplasmic and nuclear extracts (Figure 5e). Cytoplasmic partners of mTOR, Raptor and Rictor, robustly precipitated with mTOR in cytoplasmic extracts (Figure 5e). In nuclear extracts, we could confirm mTOR-RNA Pol2 and -Ell interactions, whereas interactions with Cdk9 and Paf1 were not detected in western blot assays. We conclude that nuclear mTOR associates with or is positioned in the vicinity of RNA Pol2 complexes. Both initiating (Ser5-phosphorylated) and elongating (Ser2-phosphorylated) RNA Pol2 co-precipitated with nuclear mTOR.

### mTOR-bound genes show high levels of polymerase pausing

Developmental promoters are kept repressed but promptly activatable thanks to the dynamic antagonism between PRC and RNA Pol2 complexes^17,36^. PRC1 was previously shown to counteract and disassemble the initiating polymerase. Despite this, initiating RNA Pol2 is abundant at developmental gene promoters^17,36^. Having detected RNA Pol2 as a co-precipitate of mTOR in the nucleus, we next investigated whether mTOR+ PRC+ targets show distinct Polymerase occupancy compared to other PRC targets. For this, we reanalyzed previously published ChIP-seq datasets for initiating, elongating, and total Pol2^16^ (Figure 6a, b). We observed that mTOR targets feature higher levels of the initiating RNA Pol2 (Ser5-phosphorylated), which span a broader area around the TSS (Figure 6a-d). Indeed, mTOR+ PRC+ gene promoters are distinct from mTOR-PRC+ genes in that these show high Pol2 Ser5p at promoters and lower Pol2 Ser2p (elongating Polymerase) at gene bodies (Figure 6d). Since this suggests TSS-proximal pausing of RNA Pol2, we calculated the Polymerase pausing index (PI) for each gene and found that mTOR+ PRC+ genes show significantly higher pausing compared to mTOR-PRC+ genes (Figure 5e, S5c-e). Thus, mTOR binding distinguished expression-poised early developmental regulators from more actively transcribed targets of the Polycomb machinery and may mediate their timed activation during development.

**Figure 6.**
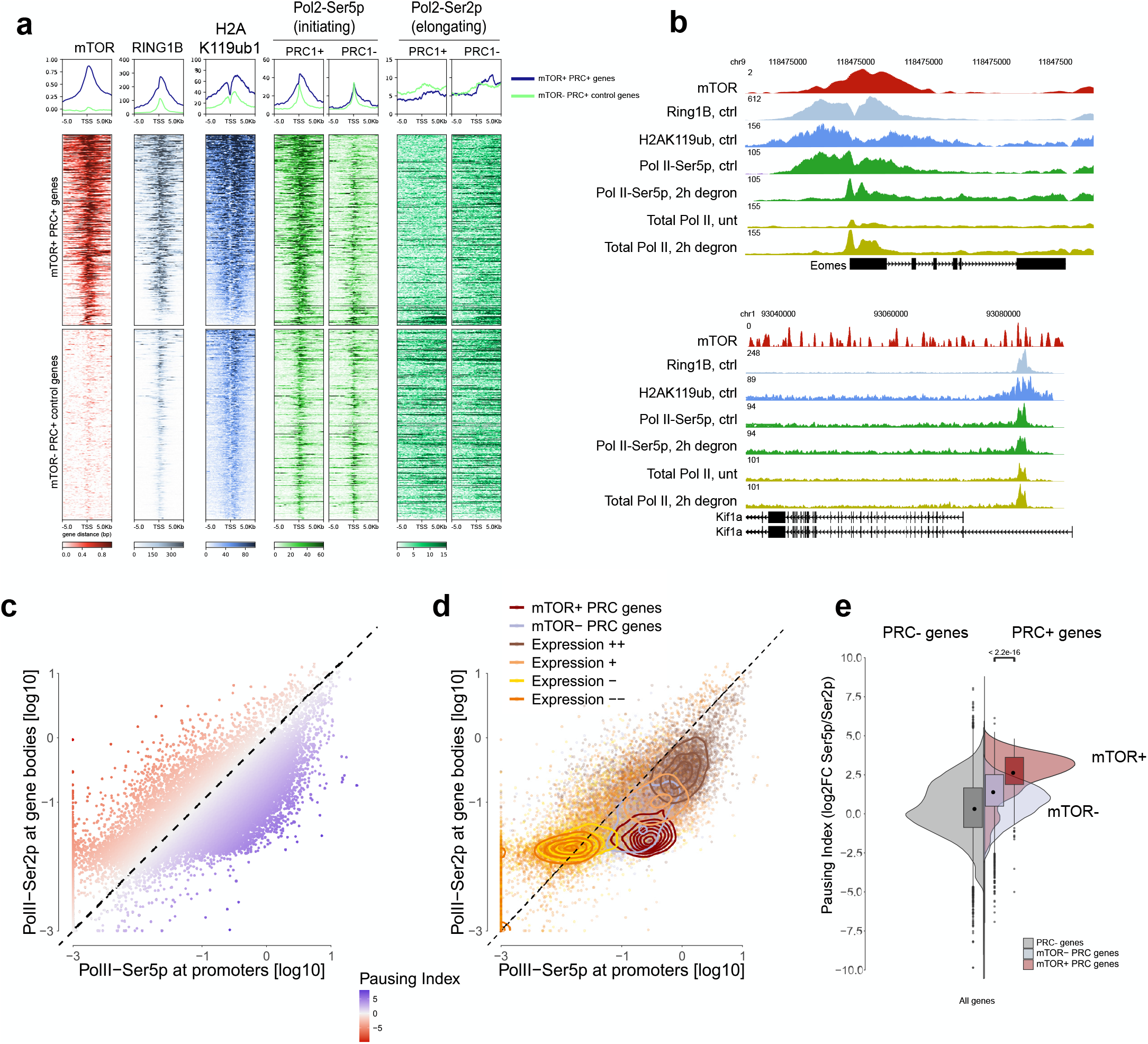
mTOR binds at developmental genes with high levels of Pol2 pausing at promoters. a. Heatmap showing levels of mTOR, Ring1b, H2AK119ub1, RNA Pol2-Ser5p and -Ser2p +/-5kb of the TSS of at mTOR+PRC+ genes vs mTOR-PRC+ control genes. b. Genome browser snapshot of a representative mTOR+PRC+_ gene (top) and mTOR-PRC+ gene (bottom). c. Scatter plot showing average levels of Pol2-Ser2p at gene bodies and Pol2-Ser5p at promoters at all genes. Color denotes Polymerase pausing index (PI). For details see Figure S5c. d. Same plot as in (c), with mTOR+ PRC+ genes and mTOR-PRC+ genes colored distinctly. PRC-genes are colored according to expression quantiles (see Figure S5d, e). Lines denote density. mTOR+ PRC+ genes show higher pausing indices compared to genes of comparable expression levels. e. Violin plots showing pausing indices at the shown gene groups.

## DISCUSSION

Since its discovery in the 1990s, mTOR has been shown to command major cellular processes such as protein synthesis and metabolism^19^ and has also been implicated in transcriptional control^22,23,38^. Despite ubiquitous knowledge about its activities across tissues, and maybe because of it, mTOR has by and large been neglected in pluripotency regulation. Here we identify a divergent new role for mTOR in stage-specific gene regulation in pluripotent ESCs.

We have previously shown that inhibition of mTOR pauses embryonic development by putting blastocyst-stage embryos and ESCs in a diapause-like dormant state^29^. In addition, *mTOR* knockout embryos fail to develop beyond E5.5^25^. Perturbation of mTOR-controlled processes such as protein synthesis and autophagy perturb development, however at earlier or later stages, and do not replicate the mTOR loss of function phenotypes in either case^26,27,39,40^.

The role of Polycomb in embryonic development has been known at the functional level for over 80 years^41^. The molecular circuits were revealed by contributions from many groups over the last decades^42^. Among PRC1-targets, master regulators of development make up the most heavily repressed gene set, with other genes (e.g. metabolic) experiencing lighter repression, leading to higher transcription levels^18,43^. Despite higher repression, developmental regulators react faster and stronger to the lifting of PRC repression, revealing their readiness to be activated. The presence of initiating Polymerase II is likely an essential player for this acute responsiveness. We propose that mTOR synergizes with the Polycomb and Pol2 machineries for correctly-timed activation of master developmental regulators.

In mouse ESCs, a subset of PRC1-target developmental genes was shown to be more firmly repressed in S and G2 phases of the cell cycle compared to G1^44^. Along the same line, ESCs in the G1 phase more efficiently differentiate^45,46^. ESC growth and proliferation rate is controlled by mTOR, in addition to other pathways such as Wnt^2,25,29,47^. mTOR’s dual role in regulating cell growth and developmental genes, may underlie the distinct propensity for differentiation in the G1 phase. Although it remains open whether chromatin-engaged mTOR assumes a transcription-activating or - inhibitory role, we suspect that its activity and distribution across different cellular domains is a key player in timing of developmental decisions.

mTOR inhibition pauses mouse development in vitro by putting embryos into a diapause-like state^29^. Our evidence suggests that while inhibition of cytoplasmic mTOR activities rewires cellular anabolism for energy preservation^30^, inhibition of nuclear mTOR may stabilize pluripotency. It has been shown that the uterine fluid is deprived of nutrients and growth factors during diapause (or vice versa, the deprivation of nutrients may initiate diapause)^48,49^ and that these rebound at the time of diapause exit. The increased availability of nutrients and growth factors reactivates mTOR^44^. Together with the resumption of cellular growth via cytoplasmic mTOR activity, these permissive signals may reenrich for nuclear mTOR at its chromatin targets.

mTOR does not carry a canonical nuclear localization signal and there are currently no known nuclear chaperones or importers into the nucleus. Few residues have been implicated in nuclear-cytoplasmic shuttling control^21^. We have generated ESC lines with mutations in these residues however these did not result in a significant change in mTOR localization (data not shown). Furthermore, mTOR does not contain a DNA-binding domain nor is it clear how mTOR recognizes H2AK119ub1 at its target promoters. Thus, at least in mESCs, the full recruitment and binding process of nuclear mTOR remains to be elucidated.

Taken together, our results thus point to nuclear mTOR as a chromatin regulator in pluripotent cells and suggest that it may link information of the cell’s microenvironment to gene expression programs.

## ACKNOWLEDGMENTS

We thank members of the Bulut-Karslioglu Lab for discussions and feedback, scientific service facilities of the Max Planck Institute for Molecular Genetics, especially the flow cytometry, microscopy and sequencing cores, for excellent service and discussions; Heike Stephanowitz for technical assistance in MS sample preparation. We thank Prof. Tatsuya Maeda for plasmids containing wild type and hyperactive mTOR, Prof. Jie Chen for plasmids with mutated shuttling sequences. This project was supported by the Max Planck Society (MS, PAO, C-YC, AG, DH, and AB-K), the Wellcome Trust (209400/Z/17/Z to RJK), the Humboldt Foundation (Sofja Kovalevskaja Award to AB-K), the EMBO (Young Investigator Program to AB-K), and the European Union (ERC Starting Grant, DOR CODE, 101117421 to AB-K).

## MATERIALS AND METHODS

### Cell lines and culture conditions

#### Generation of mTOR iKO ESCs

LoxP sites were inserted upstream and downstream of the mTOR locus using the CRIS-PITCh strategy^51^. The PITCh donor plasmid, containing loxP and either puromycin or hygromycin (for downstream or upstream targets, respectively) with 120 bp flanking homology, was cloned into the pCRIS-PITChv2-FBL vector. sgRNAs were designed using the Benchling gRNA design tool. sgRNAs for both upstream (GCCAGGCAAGTGTTAATGCG) and downstream (ggaaaaacccaatcctaacg) target sites were cloned into the px330A 1×2 plasmid. Golden Gate assembly was used with px330A 1×2-target gRNA and px330S-2-PITCh-gRNA plasmids to generate an all-in-one CRISPR-Cas9 plasmid.

The pCRIS-PITChv2–downstream loxP and Cas9 vectors were co-nucleofected on E14 mESCs line using the Amaxa 4D Nucleofector Kit (Lonza V4XP-3024). After a one-day recovery, cells were selected using puromycin (0.5 µg/ml) for 5 days. Single colonies were picked into 96-well plates for genotyping. Similar methods were used for the sequential knock-in of the upstream loxP site.

The knock out was induced by nucleofection with Cre-GFP (4 µg) plasmid per 1 million mTOR^loxp/loxp^ mESCs using the Lonza 4D nucleofector kit. The cells were sorted for CRE-GFP +ve 48 hours after nucleofection for experiments. The following primers were used for genotyping PCRs.

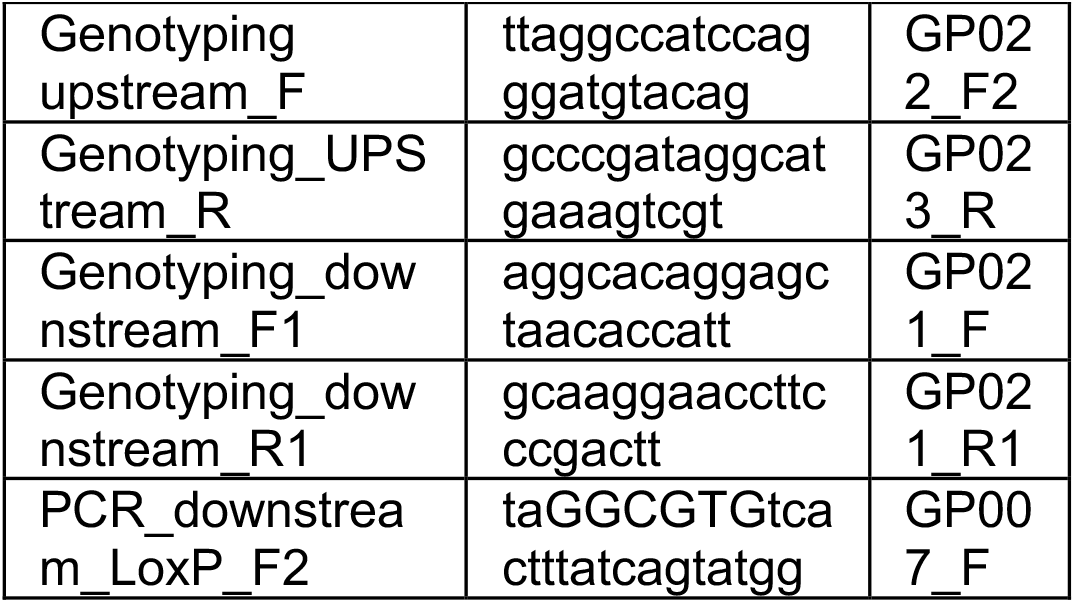
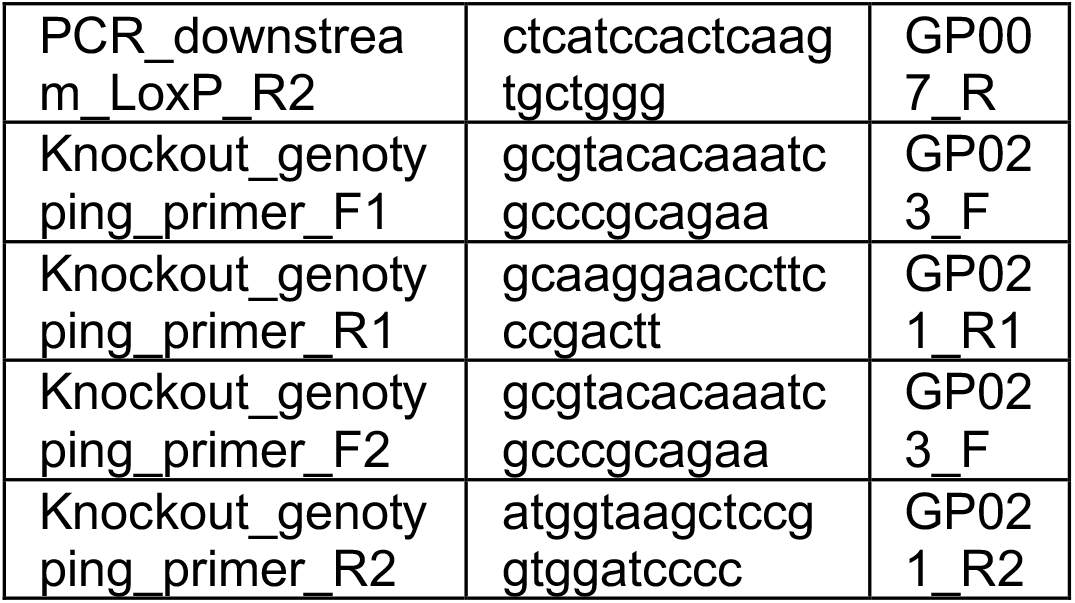

#### Generation of mTOR-dTAG ESCs

The design and cloning strategy of endogenous tagged FKBPF36V-mTOR mESCs using the CRISPR-PITCh system^51^. A gRNA targeting the first mTOR coding exon (exon 2) was designed using Benchling (sequence CACCGTCAGGGCAAGATGCTTGGGA). gRNA was cloned into the pX330A CRIS-PITCh vector as above. Primers to amplify the knock-in FKBPF36V cassette from pCRIS-PITChv2-Puro-dTAG(BRD4) (Addgene #91793) were designed to incorporate microhomologies for mouse MTOR (exon 2). Since the cut site introduced by mTOR sgRNA is 4 bp downstream to the transcription start site, frame shift correction was taken into consideration when designing the primers. The cassette PCR was performed using Q5 Hot Start High-Fidelity 2X Master Mix (NEB, M0494L). PCR products were resolved on an agarose gel and the band corresponding to 1215 bp was purified using QIAquick Gel Extraction Kit (Qiagen, 28706). The knock-in FKBPF36V cassette was assembled with pCRIS-PITChv2-Puro-dTAG(BRD4) (MluI-HF digested (NEB, R3198S) using NEBuilder HiFi DNA Assembly Master Mix. The assembly products were used to transform *E. coli* (*DH5a*). Single colonies were picked and the sequence of pCRIS-PITChv2-Puro-dTAG(MTOR) was confirmed using Sanger sequencing (with primers dTAG-seq-FWD/REV). pX330A-Mtor/PITCh and pCRIS-PITChv2-Puro-dTAG(MTOR) were co-transfected into mESCs using an Amaxa 4D Nucleofector X Unit (Lonza) according to the manufacturer’s instructions. Two days after nucleofection, the mESCs were selected using puromycin (Sigma, A1113803) for at least 5 days. Single mESC clones surviving puromycin selection were picked and expanded to establish monoclonal cell lines. To screen for copies of insertion, the clones were genotyped using PCR. For clones with positive insertion the sequence was further confirmed.

To test mTOR degradation after dTAG-13 treatment, mESCs were treated with 500 nM dTAG-13 (Torcis, 6605) and subjected to protein extraction and western blot analysis.

#### ESCs in serum/LIF media

E14 mouse ESCs were cultured without feeders on 0.1% gelatin-coated dishes (Sigma-Aldrich, G1393) with daily media change and were passaged every other day. Culture media contains DMEM/High glucose with Glutamax (Thermo, 31966047), 15% FBS (Thermo, 2206648RP), 1x NEAA (Gibco, 11140-035), 1x Penicillin/streptomycin (Life Technologies, 15140148), 0.2% β-mercaptoethanol (Thermo, 21985023) and 1000 U/mL LIF (homemade). Cells were cultured at 37°C under 20% O_2_ and 5% CO_2_. Cells routinely tested negative mycoplasma.

#### ESCs in 2i/LIF media

N2B27 media (1:1 neurobasal media (Thermo, 21103-049) and DMEM/High glucose with Glutamax media, 1x NEAA, 1x Penicillin/streptomycin, 1x Glutamax (Thermo, 61870044), 15% BSA fraction V (Gibco, 15260-037), 0.2% β-mercaptoethanol (Thermo, 21985023)) was supplemented with 1 µM PD0325901 (Tebubio, 25704-0006), 3 µM CHIR-99021 (Sigma, SML1046) and 1000U/mL LIF.

#### Mouse embryonic fibroblasts (MEFs)

MEFs were cultured in media containing DMEM/High glucose with Glutamax (Thermo, 31966047), 10% FBS (Thermo, 2206648RP) and 1x Penicillin/streptomycin (Life Technologies, 15140148).

### Chemical treatments and differentiation

INK-128 (MedChemExpress, HY-13328, 200 nM final), RapaLink-1 (MedChemExpress, HY-111373, 200 nM final), and cycloheximide (Biomol, 54646.1, 100 nM final) were used. To induce differentiation, cells were washed once in PBS before being supplemented with fresh media without LIF.

### SSEA1-flow cytometry

Wild-type ESCs (control or mTORi) were dissociated from plates using TrypLE (Thermo Fisher 12604-021) or mTOR iKO (CRE-GFP +ve) cells collected from FACS sorting were washed in Dulbecco’s phosphate-buffered saline (Dulbecco). Cells were labeled with Alexa488-SSEA1 (BioLegend, 125610, 1:1,000) or Alexa647-SSEA1 (BioLegend, 125607, 1:500) for 20 min in the dark on ice and subsequently washed in phosphate-buffered saline (PBS) containing 2% bovine serum albumin (BSA). After washing, fluorescence was measured on a BD FACSAria Fusion or BD FACSAriaII flow cytometer. Data analysis and visualization was performed using FlowJo (v10.8.2).

### Alkaline phosphatase staining

Alkaline phosphatase staining was done using Vector Red Substrate Kit (Vector Laboratories, VEC-SK-5100) following manufacturer’s guideline. Cells were washed once in PBS and incubated with the freshly made substrate working solution (For 5 ml of 100 mM Tris-HCl, pH 8.2, added 2 drops of Vector Red Reagent 1, 2 drops of Vector Red Reagent 2 and 2 drops of Vector Red Reagent 3) at incubated at 37 °C for 20 min. The substrate working solution was removed and the cells were washed in PBS for 10 min before imaging. Bright field images were taken by an Olympus CKX53 microscope.

### Cell counting and shape analyses

For colony counting and morphological analysis, Cell Profiler^52^ v4.2.8 was used. Bright field images were used as input and were converted to grayscale. Primary objects were identified using an adaptive thresholding strategy (minimum cross-entropy) in the size range of 75-300 pixel units. Morphological measurements were extracted using the ObjectSizeShape module and plotted using GraphPad Prism v10.

### Immunofluorescence imaging

Cells were cultured on gelatin-coated glass slides as colonies and subsequently fixed with 4% PFA in DPBS. After fixation cells were washed 3 times and permeabilized using PBS-0.2% Triton X-100 for 10 min at RT. After permeabilization cells were blocked in DPBS-0.2% Triton X-100 containing 2% BSA and 5% goat serum for 1h at RT. After blocking, cells were incubated with primary antibodies in the blocking buffer (1:200 NANOG (Abcam, ab80892), 1:500 SOX2 (Abcam, ab79351) overnight at 4℃ and subsequently washed three times in PBS-0.2% Triton X-100 containing 2% BSA at RT. After washing, cells were incubated with secondary antibodies in blocking buffer (1:700 Donkey anti-Rabbit IgG (H+L) Alexa Fluor™ 568 (Thermo Fisher Scientific, A10042), 1:700 Donkey anti-Mouse IgG (H+L) Alexa Fluor™ 488 (Thermo Fisher Scientific, A21202)) for 1h at RT. Subsequently, the cells were washed in PBS-0.2% Triton X-100 containing 2% BSA at RT. Cells were embedded in Vectashield containing DAPI (Vectashield, Cat: H-1200) sealed with a cover glass (Brand, 470820). Images were acquired with ZEISS LSM880 microscope at 20x magnification. Images were further processed using Fiji (version 2.3.0).

### Generation of mTOR overexpression ESCs

Gene cassettes containing mCherry, P2A, FLAG-tag or myc-tag and mTOR variants were cloned into a pSLQ2818 pPB plasmid (Addgene, #84241), where the sequences between the 3’- and 5’-ITR were replaced. The FLAG-mTOR^WT^ and FLAG-mTOR^hyper^ were amplified from pcDNA3.1-FLAG-mTOR and pcDNA3.1-FLAG-mTOR^SL1+IT^ respectively. The myc-mTOR^KD^ was amplified from a myc-mTOR kinase-dead plasmid (Addgene, #8482). The PiggyBac plasmids were co-transfected into ESCs using an Amaxa 4D Nucleofector X Unit (Lonza) according to the manufacturer’s instructions. To screen for positive integrations, the cells were sorted 3 days after transfection for mCherry fluorescence using a FACS Aria II flow cytometry, followed by a second sorting after another 3 days.

### Protein extraction and western blotting

#### Subcellular fractionation of cytoplasm, nucleus and chromatin

5×10^6^ cells were pelleted, washed once in ice-cold PBS, and resuspended in 500 µl cytoplasmic lysis buffer (10 mM HEPES pH 7.9, 10 mM KCl, 340 mM sucrose, 10% glycerol, 1 mM DTT, 0.1% Triton X-100, 1x protease and phosphatase inhibitor cocktail, 1 mM PMSF, 5 mM NaF, 1 mM Na_3_VO_4_) on ice for 10 min. Lysates were centrifuged at 1300 x g for 5 min at 4°C and cytoplasmic fraction was collected. Nuclear pellet was washed once in cytoplasmic lysis buffer, spun at 1300 x g for 5 min at 4°C and resuspended in 500 µl of nuclear lysis buffer (3 mM EDTA, 0.2 mM EGTA, 1 mM DTT, 1x protease and phosphatase inhibitor cocktail, 1 mM PMSF, 5 mM NaF, 1 mM Na_3_VO_4_) and incubated on ice for 5 min. Lysates were centrifuged at 1700 x g for 5 min at 4°C and nucleoplasmic fraction was collected. Chromatin pellet was resuspended in 100 µl 1x Laemmli Sample Buffer (Bio-Rad, 161-0747) containing 5% β-mercaptoethanol and sonicated using a Bioruptor for 5 cycles (30 sec ON, 30 sec OFF). Finally, the cytoplasmic and nucleoplasmic fractions were mixed with 1x Laemmli Sample Buffer and 5% β-mercaptoethanol and together with the chromatin fraction samples boiled at 99 °C for 5 minutes.

#### SDS-PAGE and western blotting

The denatured samples were loaded on a 4-15% Mini-PROTEAN TGS precast protein gel (Bio-Rad, 4561083) and separated by electrophoresis at 200 V for 30 min using 1x Tris/Glycine/SDS running buffer (Bio-Rad, 1610772). Proteins were transferred to a PVDF membrane (Thermo Fisher IB24001) using the pre-programmed method P0 on an iBlot2 gel transfer device (Thermo Fisher IB21001). Membranes were incubated first in blocking buffer (5% non-fat milk in 1x TBST (Thermo Fisher, 28360)), then in specific primary antibodies diluted in blocking buffer (mTOR 1:1000 (Cell Signaling Technology, 2983), vinculin 1:1000 (Sigma, V9131), β-actin 1:2000 (Abbkine, A01010), H2A 1:1000 (Cell Signaling Technology, 12349), pS6 1:1000 (Cell Signaling Technology, 4858)) at 4°C overnight. Membranes were washed 3 times in 1x TBST for 30 min before incubation with secondary antibodies (mouse anti-rabbit IgG 1:2000 (Jackson Immuno Research, 211-032-171) or goat anti-mouse IgG 1:5000 (Jackson Immuno Research, 115-035-174) at room temperature for 1h. After washing, signal was developed using SuperSignal WestDura duration substrate (Thermo Fisher, 34075). The blots were imaged on a ChemiDoc imaging system (Bio-Rad).

### ChIP-seq

After washing once in PBS, ESCs were incubated with 1% formaldehyde (Thermo Fisher Scientific, 28906) at room temperature for 10 min. The fixative was quenched with 125 mM glycine for 5 minutes. Cells were then washed 2 times in ice-cold PBS before incubation at 4 °C in swelling buffer (25 mM HEPES, pH 7.9, 1.5 mM MgCl_2_, 10 mM KCl, 0.1% Igepal 630, 1x protease and phosphatase inhibitor cocktail (Thermo Fisher, 78443), 1 mM PMSF, 1 mM NaVO_3_, 5 mM NaF) for 10 min. To collect the nuclei, cells were scraped on ice, passed through a 18G needle for 5 times and spun at 3000 x g for 5 minutes at 4°C. Nuclei pellets were resuspended in 1 ml of sonication buffer (50 mM HEPES, pH 7.9, 140 mM NaCl, 1 mM EDTA, 1% Triton X-100, 0.1% sodium deoxycholate (Thermo Fisher Scientific, 89904), 0.1% SDS, 1 mM PMSF, 1 mM NaVO3, 5 mM NaF) and incubated on ice for 10 min. Resulting chromatin was sheared to an average size of 500-1000 bp using an E220 Evolution Covaris sonicator for 6 cycles (1 minute per cycle). Chromatin immunoprecipitation was performed by mixing chromatin containing 25 µg of DNA, 1 µg of mTOR antibody (Abcam, ab32028) and 20 µl Protein A dynabeads (Thermo Fisher Scientific, 10002D, pre-washed and resuspended in sonication buffer) and incubated at 4°C overnight. For 1% input control, 0.25 µg of chromatin was set aside in 100 µl sonication buffer at 4°C. The beads were washed twice in sonication buffer, once in high salt wash buffer (sonication buffer with 500 mM NaCl instead) and once in TE buffer (10 mM Tris-HCl, pH 8.5, 1 mM EDTA, 1% SDS). After washing steps, the beads were resuspended in 100 µl elution buffer (50 mM Tris-HCl, pH 7.5, 1 mM EDTA, 1% SDS). 1 µl of RNaseA (Thermo Fisher Scientific, EN0531) was added to the samples and incubated at 37°C for 30 min with shaking, followed by incubation with 5 µl of Proteinase K (NEB, P8107S) and incubated at 65 °C for 2 hours with vigorous shaking. DNA was purified using ChIP DNA Clean & Concentrator columns (D5205, Zymo Research) and the concentration was measured using Qubit dsDNA HS Assay (Invitrogen, Q32851). Sequencing libraries were prepared using the NEBNext Ultra II DNA Library Prep Kit (NEB, E7645L) according to manufacturer’s instructions. Sequencing libraries were sequenced on a NovaSeq 6000 in 100-bp paired-end mode and 50 million fragments per library.

### Generation of the *Ring1a/1b*^*dTAG*^ cell line

To allow for their rapid depletion, we introduced an N-terminal dTAG into the endogenous *Ring1a* and *Ring1b* genes using CRISPR/Cas9-mediated genome editing. sgRNAs specific for the 5’ end of each gene were designed using the CRISPOR online tool (http://crispor.tefor.net; Ring1a guide = tgcattcgccggcgtcgtca, Ring1b guide = tcaaccattaagcaaaacat) and oligonucleotides encoding sgRNAs were cloned into the pSpCas9 (BB)-2A-Puro plasmid (Addgene #62988). Targeting constructs containing the FKBP12F36V tag (dTAG) coding sequence (obtained from Addgene plasmid #91797) flanked by 700-1000 bp homology arms were cloned by Gibson Assembly (NEB). E14 ESCs were transfected at approximately 70% confluency in a 6-well plate with 0.5 µg of each Cas9 guide plasmid and 2 µg of each targeting construct using Lipofectamine 3000, according to the manufacturer’s protocol (Thermo Fisher Scientific). The following morning, transfected cells were passaged onto new plates at a range of low densities and selected with 1 µg/ml puromycin for 48 hours. Approximately 7 days later, individual colonies were picked into 96-well plates. Genomic DNA samples from individual clones were subsequently used for a PCR screen to identify clones that were homozygous for dTAG insertion at both *Ring1a* and *Ring1b*. Putative Ring1a/1b^dTAG^ lines were further expanded and subjected to treatment with 100 nM dTAG13 (Tocris) for 2 hours. Cellular extracts were then analysed by western blot using antibodies against RING1B (Cell Signaling Technology #5694) and H2AK119ub1 (Cell Signaling Technology #8240).

### Induction of PRC1 catalytic mutation

The E14 ESC PRC1^CPM^ line was generated previously by engineering a targeting construct comprising exon 3 of *Ring1b* in forward orientation (flanked by 100 bp of *Ring1b* intron 2/intron 3) followed by a mutant copy of exon 3 (encoding I53A and D56K mutations) in reverse orientation (flanked by splice donor and acceptor sites from mouse *IgE* gene). CreERT2 was additionally inserted into the *Rosa26* locus. To induce the catalytic mutation via exon inversion, cells were treated with 800 nM 4-OHT for the indicated time periods. During this time frame, H2AK119ub1 was lost, while parental PRC complex remained intact.

### CUT&Tag

Cleavage Under Targets and Tagmentation (CUT&Tag) was performed on freshly isolated nuclei or cells based on Kaya-Okur et al. with minor modifications^53^. Two biological replicates were performed.

For freshly isolated nuclei: For each CUT&Tag reaction, 3.5 µl of Concanavalin A beads (Polysciences, 86057) were equilibrated by washing 2 times in 100 μl Binding buffer (20 mM HEPES-KOH, pH 7.5, 10 mM KCl, 1 mM CaCl2, 1 mM MnCl2) and concentrated again in 3.5 μl Binding buffer. Beads were added to 1×10^5^ cells and were incubated for 10 min with rotation at room temperature. The nuclei-bound beads were separated on a magnet and resuspended in 25 μl cold Antibody buffer (Wash buffer with 0.1% BSA) containing diluted primary antibody or IgG (H3K27me3 1:100 (Cell Signaling Technology, 9733, H2AK119ub1 1:100 (Cell Signaling Technology, 8240), IgG 1:100 (Abcam, ab46540)). Samples were incubated at 4°C with gentle nutation overnight, washed, and incubated with matching secondary antibody (guinea pig α-rabbit antibody, 1:100 (Antibodies online, ABIN101961) and were incubated for 30 min at room temperature.

For tagmentation, 3xFLAG-pA-Tn5 (homemade) preloaded with mosaic-end adapters was diluted 1:250 in wash buffer (20 mM HEPES-KOH, pH 7.5, 300 mM NaCl, 0.5 mM Spermidine, 1 mM PMSF, 5 mM NaF, 1 mM Na_3_VO_4_, 1x protease phosphatase inhibitor cocktail) and 50 µl of it was used to resuspend the beads. Beads were incubated for 1 hour at room temperature with gentle nutation and were washed once in 200 µl of wash buffer on a magnet. Tagmentation was performed by incubating the beads in 50 µl Tagmentation buffer (10mM MgCl2 in 300-wash buffer) for 1 hour at 37 ℃. Tagmentation was stopped by adding 2.25 µl 500 mM EDTA and 2.75 µl 10% SDS to the beads followed by adding 0.5 µl of Proteinase K (20 mg/ml, Invitrogen, AM2546). After vortexing for 5 sec, beads were incubated at 55°C for 1 hour and then at 70°C for 30 minutes. Resulting DNA was purified using ChIP DNA Clean & Concentrator columns (D5205, Zymo Research). DNA was amplified using NEBNext HiFi 2x PCR Master Mix (New England BioLabs, M0541L) and i5- and i7-unique barcoded primers for 8 cycles. Ampure XP beads (Beckman Coulter, A63881) were used for post-PCR cleanup. Library quality was assessed using Agilent High Sensitivity D5000 ScreenTape System and Qubit dsDNA HS Assay (Invitrogen, Q32851). Sequencing libraries were pooled in an equimolar ratio and sequenced on a NovaSeq 6000 in 100-bp paired-end mode and 5-8 million fragments per library.

### Computational analysis

The computational analysis that has not been conducted by the use of tools was done in R 4.2.26 (R Core Team (2022). R: A language and environment for statistical computing. R Foundation for Statistical Computing, Vienna, Austria. URL https://www.R-project.org/). If not stated otherwise, the generated plots were generated using ggplot2^54^ 3.4.4.

#### CUT&Tag

The reads were processed partially according to the pipeline described by Zheng et al^55^. First, the reads of the samples were trimmed using Trim Galore/cutadapt 2.417^56^. The trimmed reads were aligned to the mm10 genome using bowtie2^57^ 2.5.018 with parameters *--end-to-end --very-sensitive --no-mixed --no-discordant --phred33 -I 10 -X 700*. The resulting SAM files were filtered (mapping quality >= 2) and converted to BAM files using samtools^58^ view 1.17. The BAM files were sorted and indexed using sambamba^59^ sort 0.8.2. For spike-In calibration, the trimmed reads were additionally mapped to the E. coli genome using bowtie2 with parameters settings *--local --very-sensitive --no-overlap --no-dovetail --no-mixed --no-discordant --phred33 -I 10 -X 700*.

Scaling factors were computed according to the SRPMC normalization method as described by DeBerardine (2022)^57^ :

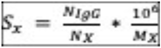

With 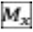 being the number of reads in sample 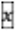 that mapped onto the mm10 genome.

Calibrated BigWig coverage files were generated from the BAM files using deepTools’ bamCoverage 3.5.4.post1 with a smoothing window of 300bp and the before-mentioned scaling factors. The coverage files of replicates were merged by taking their mean using deepTools’ bigwigCompare 3.5.4.post1 with pseudo-count 0 and operation “mean”.

### ChIP-seq

The ChIP-seq samples were first trimmed with Trim Galore and then mapped to the mm10 genome using bowtie2 with default parameters. The SAM files were converted to BAM files using samtools view. The BAM files were sorted and indexed using sambamba sort. Unmapped and duplicate reads were removed from the BAM files using sambamba view such that only uniquely mapped reads remain.

Furthermore, smoothed (300bp windows) and CPM normalized BigWig files were generated using deepTools’ bamCoverage. The resulting CPM normalized coverage files were further normalized for background by forming the log2 ratio of the mTOR and input coverage tracks using deepTools’ bigwigCompare (*--pseudocount 1 --operation log2*).

Finally, the coverage files of replicates were merged by taking their mean using deepTools’ bigwigCompare (*--pseudocount 0 --operation mean*).

For calling mTOR peaks, MACS2^57^ peakcall 2.2.7.125 was used and the broad mode (*--broad*) with default parameters. The peaks for the 2i and MEF mTOR ChIP-seq samples were called without an input experiment. For the remaining samples input experiments were used as control.

#### mTOR Target Gene Identification and Analysis

Prior to identifying target genes of mTOR,any peaks overlapping with blacklisted regions were removed from the samples.

To find target genes, the TSS of the mouse genes were extracted using either TxDb.Mmusculus.UCSC.mm10.knownGene (<u>10.18129/B9.bioc.TxDb.Mmusculus.UCSC.mm 10.knownGene</u>) v3.10.07 (for mm10 genome) from Bioconductor^60^ v1.30.209. Peaks located 2kb up- or downstream of a TSS were deemed as targeting the respective genes. Genes targeted by mTOR in all ChIP-seq replicates of mTOR were considered an mTOR ESC target gene.

To cluster the mTOR target genes, deepTools’ compute Matrix reference-point (*--referencePoint center -b 1000 -a 1000*) was used to compute the profiles of the normalized coverage of H2AK119ub1 in ESC 1kb up- and downstream from the TSS of the mTOR target genes. The coverage levels were summed up per gene over all bins log10-transformed. The resulting values were centered around their mean and scaled based on their standard deviation and then used as input for k-means clustering. The clustering was performed using the function kmeans (2 clusters, *nstart = 50*) of the R package stats 4.2.2 with seed 123.

Gene ontology analysis was carried out using the function compareCluster of the R package clusterProfiler^61^ v4.6.010 with a cut off of 0.05 for p-values and 0.1 for q-values. The p-values were corrected using the Benjamini-Hochberg procedure. The gene set enrichment analysis based on H2AK119ub1 levels was performed using the function gseGO of the clusterProfiler package. Prior to running the function, the H2AK119ub1 levels (computed as described before) were log-scaled. Then, the log2 FC of the values vs. the mean was computed and the genes were ordered according to them. Based on those values, the gene set enrichment analysis was performed.

#### PRC Target Gene and Control Gene Identification

For the identification of Polycomb target genes, reads from Suz12 and Ring1B ChIP-seq experiments were first merged by replicate. The individual replicates were processed and peaks were called in the same way as described for the mTOR ChIP-seq samples. For every replicate, target genes were identified based on the same criterion as for the mTOR samples (+/-2kb from gene TSS). PRC1 target genes were defined as genes identified in all three Ring1B replicates and PRC2 targets as genes identified in all three Suz12 replicates. PRC targets were then identified by intersecting PRC1 and PRC2 target genes. Furthermore, genes identified as PRC targets and mTOR targets were considered mTOR+ PRC targets and genes only identified as PRC targets were considered mTOR-PRC targets. To analyse the importance of H2AK119ub levels, mTOR-PRC targets were categorized into four quantiles based on their H2AK119ub levels in WT conditions, Q1 containing the targets with the highest levels and Q4 those with the lowest levels.

Since mTOR targets are defined using the overlap of 5 samples in total, a high number of PRC targets were categorised as mTOR-PRC targets even though mTOR peaks were called around their promoters in at least one replicate. To ensure a high confidence set of mTOR negative PRC targets as a basis for control genes, mTOR-free PRC targets were defined by filtering all mTOR-PRC targets that have mTOR peaks in at least one replicate. Then, mTOR-free targets in the Q1 (highly ubiquitinated targets) PRC target group were extracted and further downsampled to the same number as mTOR+ PRC targets (n = 312). Subsequently, this sub-sample of Q1, mTOR-free samples was used as control genes.

#### Pausing Index analysis

To assess pausing of the Polymerase, the published ChIP-seq coverage files of the Ser2/Ser5-phosphorylated RNA-Polymerase II were used as published by Dobrinić et al. (2021). To ensure similar scaling, the Ser2p- and Ser5p-RNA Pol II tracks were first quantile normalized condition-wise (untreated and 2h Auxin induction) using the r function normalize.quantiles. Then, the TSS region (TTSR) is defined as 50bp upstream and 300bp downstream of the TSS and the gene body region (GBR) is defined as 300bp downstream from the TSS to 3kp downstream from the TES (coordinates as extracted from TxDb.Mmusculus.UCSC.mm10.knownGene).

The quantile normalized coverage tracks are used to compute mean Ser5p-RNA Pol II enrichment in the TSSR and mean Ser2p-RNA Poll II enrichment in the GBR. These values were subsequently used for visualisation in scatter plots using ggplot and to compute the Pausing Index (PI) as the log2 fold change of mean Ser5p enrichment in TSSR and mean Ser2p enrichment in GBR. The PI definition was taken from Day, D.S., Zhang, B., Stevens, S.M. et al. (2016)^50^, but adjusted to use Ser5p- and Ser2p-RNA Poll II tracks instead of total RNA Pol II.

### Visualizations

#### deepTools heatmaps and profiles

deepTools’ computeMatrix reference-point (*--referencePoint center -b X -a X*) was used to compute coverage levels of the samples around Xbp up- and downstream of the TSSs of genes in the mm10 genome (usually X = 5kb). For the CUT&Tag samples, the scaled and merged BigWig files were used and for the ChIP-seq samples the input-normalized and merged BigWig files were used. The enrichment heatmaps were generated by deepTools’ plotHeatmap and the enrichment profiles by deepTools’ plotProfile.

#### Complex Heatmaps,violin plots and box plots

For the enrichment levels of the different targets, deepTools’ computeMatrix reference-point (*--referencePoint center -b 1000 -a 1000*) was used to compute coverage levels of the samples around 1kb up- and downstream of the TSSs. The values of the bins were summed up for every gene and sample resulting in a count matrix with genes as rows and the coverage samples as columns.

Negative values in the resulting count matrix were set to 0 and the final matrix was further log-scaled (pseudo-count 1) prior to visualization.

For the gene expression counts, deepTools’ multiBigwigSummary BED-file was used to compute the enrichment of the normalized BigWig files of the nucRNA-seq experiment provided by Dobrinić et al. (2021) over the gene bodies extracted from xDb.Mmusculus.UCSC.mm10.knownGene. The function computes the mean enrichment on every region which means the resulting counts were also normalized for gene length. As nuclear RNA is usually not spliced, quantification over the whole gene body (instead of only exons) is necessary. Counts were assigned to genes based on their strandedness.

Heatmaps were visualized using the function Heatmap of the R package ComplexHeatmap^62,63^ 2.15.1 and violin and box plots were generated by ggplot2.

#### Statistical tests

To assess statistical significance of the changes observed in the violin/boxplots, one-sided Wilcoxon Rank Sum tests were performed as implemented by the function wilcox.test. For the comparisons between the same genes in different conditions paired tests *(paired = TRUE, alternative = “greater”*) and for the comparisons between different gene groups within the same condition an unpaired test (*paired = F, alternative = “less”*) were performed.

#### Venn and Euler diagrams

Venn diagrams were plotted using the function venn of the R package venn. Euler plots were generated by the functions euler and plot of the R package eulerr^64^.

#### Genome browser profiles

The profiles were generated using fluff^65^. CPM normalized and merged coverage files were used to properly showcase the enrichment of mTOR over input. For the remaining figures, input-normalized coverage files were used. The normalized coverage files provided by Dobrinić et al. (2021) were used as published. For the CUT&Tag samples, the scaled and merged coverage files were used.

#### Distribution of mTOR peaks

To assess the distribution of mTOR peaks around different genomic regions, the mTOR peaks called in ESC and 0hr LIF withdrawal were intersected. Only peaks that were detected in all 5 replicates were used for the analysis. The peaks were annotated using the function annotatePeak of the R package ChIPseeker^66,67^ 1.34.1 with TxDb.Mmusculus.UCSC.mm10.knownGene as annotation file, org.Mm.eg.db as annotation data base and +/-1kb to define the TSS region. The annotation was visualized using the plotAnnoPie function of ChIPseeker.

#### Quality Control

For assessing replicated reproducibility of the datasets deepTools’ multiBigwigSummary BED-file was first used to compute mean enrichment of signal around gene promoters (TSS +/-1kb). For the ChIP data the CPM normalized tracks were used. For the CUT&Tag data the scaled (spike-in/SRPMC) tracks were used. The resulting clustering was visualized using deepTools’ plotPCA.

Mapping statistics such as sequencing depth, alignment rate, etc. were extracted from the bowtie2 log-files for the CUT&Tag data. For the ChIP-seq files, samtools view was used to count total, mapped and filtered number of read.

### Nuclear mTOR IP-mass spectrometry

#### Nuclear fractionation and IP

∼7M ESCs were harvested, washed with PBS, and resuspended in 5ml cold buffer A (10mM Hepes pH7.9, 5mM MgCl2, 0.25M sucrose, 0.1% NP40 (type Igepal630, protease inhibitors directly before use) per ∼7M cells. Cells were incubated for 10 minutes on ice, then passed 4-5 times through a 18G1 needle, and centrifuged for 10 minutes at 5000 rpm, 4C. The supernatant was transferred into a new tube - this fraction represents the cytosolic fraction.

The pellet was washed once with cold PBS, and incubated in RIPA buffer (cold) on ice for 30 min with 1x proteinase and phosphatase inhibitor (Thermo). The lysate was briefly sonicated using a biorupter in the cold room using the following parameters: intensity =low 3 cycles 30 sec on 30 sec off. For IP, Protein A/G magnetic beads were washed 3x with RIPA buffer. mTOR or IgG antibodies and washed beads were added to the lysate and incubated overnight at 4C. Afterwards, beads were washed 5 times x 10 min with RIPA buffer, then resuspended in Laemmli buffer.

#### Proteolytic digestion of immunoprecipitated proteins via SP3

Four replicates of IP’d proteins were eluted using 1X SDS-PAGE Loading dye for 10 min at 95 °C and then digested using a modified SP3 protocol^68^. Essentially, proteins were reduced with 5 mM TCEP and alkylated with 40 mM CAA for 1 h at room temperature in the dark. A 1:1 (v/v) mixture of hydrophobic and hydrophilic Sera-Mag beads was added to the sample, followed by 50% final acetonitrile (ACN). After a brief incubation the supernatants were removed and the beads were washed twice with ethanol and once with ACN. Proteins were digested in 50 mM TEAB (pH 8.5) with Lys-C and Trypsin in enzyme:protein ratio 1:200 and 1:100 wt:wt, respectively, for 16 h at 37 °C. Beads were washed twice with an excess of ACN (> 95% final). Peptides were eluted with 5% DMSO and dried until further use.

#### LC-MS of IP derived peptides

Desalted peptides from IPs were resuspended in 1% ACN with 0.05% trifluoroacetic acid (TFA) and 3 of 10 µL were injected into a Thermo Scientific Vanquish Neo system coupled on-line to an Orbitrap Exploris480 mass spectrometer (Tune version 4.2). Prior to separation, peptides were trapped on a PepMap C18 trap column (0.075 × 50 mm, 3 μm, 100 Å, Thermo Fisher). Peptide separation was performed using reverse-phase chromatography on an in-house packed C18 column (Poroshell 120 EC-C18, 2.7 μm, Agilent). Elution was carried out over a 120-minute linear gradient of increasing ACN concentration at a flow rate of 250 nL/min. Data were acquired in data-dependent acquisition (DDA) mode. MS1 scans were performed in the Orbitrap at a resolution of 120,000 over a scan range of m/z 375–1,200, with a 50% RF lens setting, 300% AGC target, and an automatic maximum injection time. MS2 scans were acquired in the Orbitrap at 15,000 resolution using a 1.6 m/z isolation window, standard AGC target, automatic maximum injection time, normalized collision energy of 30% and an automatic scan range. Only precursors with charges +2 - +4 were fragmented and dynamically excluded for 20 s. The instrument was equipped with FAIMS operated at standard settings using CVs -50 and -70 with 2 s cycle time.

#### Label-free quantification of proteins

Raw data were analyzed using FragPipe v22.0 with MSFragger v4.1, IonQuant v1.10.27^69–71^. The data were searched against a database containing the murine proteome (one protein per gene, retrieved on 2024-06-13) supplemented with common contaminants and reverse decoy sequences. The LFQ-MBR workflow in FragPipe were used with default settings if not stated otherwise, essentially: Precursor tolerance ± 10 ppm, Fragment tolerance ± 20 ppm, mass calibration and parameter optimization activated, Trypsin cleavage C-terminal of KR, 2 missed cleavages, peptide length 7 – 50, peptide mass 500 – 5,000. Enrichment and differential expression analyses were carried out using R with the limma package^72^. To this end, proteins with less than 3 out of 4 values in both conditions were removed. Data was median normalized and missing values imputed using a Perseus-like imputation.

## SUPPLEMENTARY FIGURES

**Figure S1.**
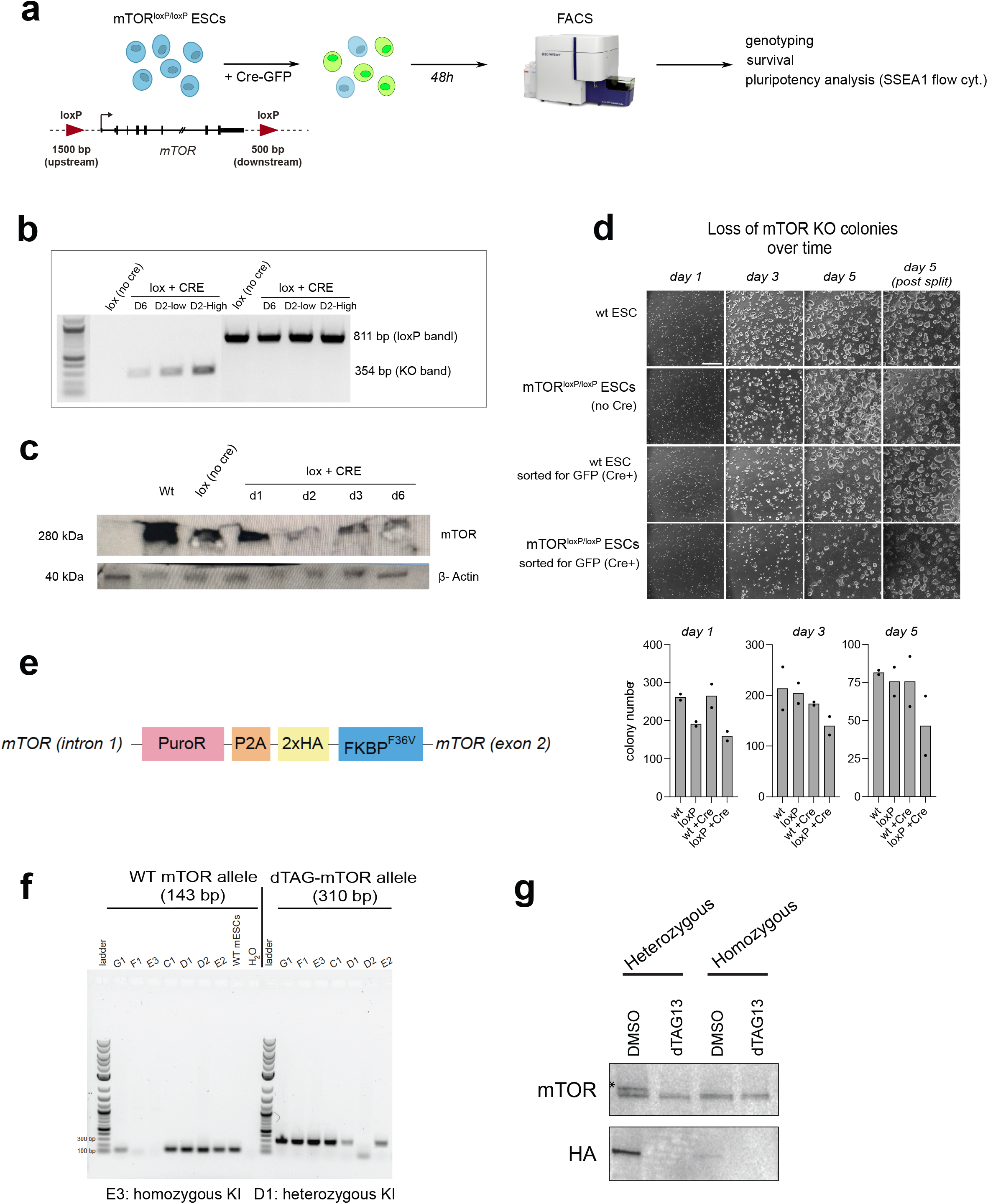
Characterization of mTOR iKO and degron ESC lines. a. Schematic showing the genetic modification and experimental workflow. b. Genotyping PCR. Two primer pairs were used, with the left set detecting deletion and right set detecting the inserted loxP (as control). Days 2 and 6 after sort are shown. c. Western blot showing levels of mTOR in wild-type ESCs, mTOR^loxP/loxP^ ESCs, and in Cre-sorted cells. Days 1-6 after sort are shown. d. Bright field pictures of ESC colonies of indicated genetic backgrounds. Bottom, number of detected colonies on respective days of culture. Colonies were counted using CellProfiler. mTOR iKO colonies are lost from culture over time, however dispersion of colonies during the split masks this effect. e. Schematic of mTOR-dTAG knock-in ESCs. f. Genotyping PCR showing the inserted dTAG allele. g. Western blot showing degradation of mTOR in the heterozygous D1 clone but not in the homozygous E3 clone. *denotes the mTOR-dTAG protein, which is not seen in the homozygous cell line despite the correct genotyping.

**Figure S2.**
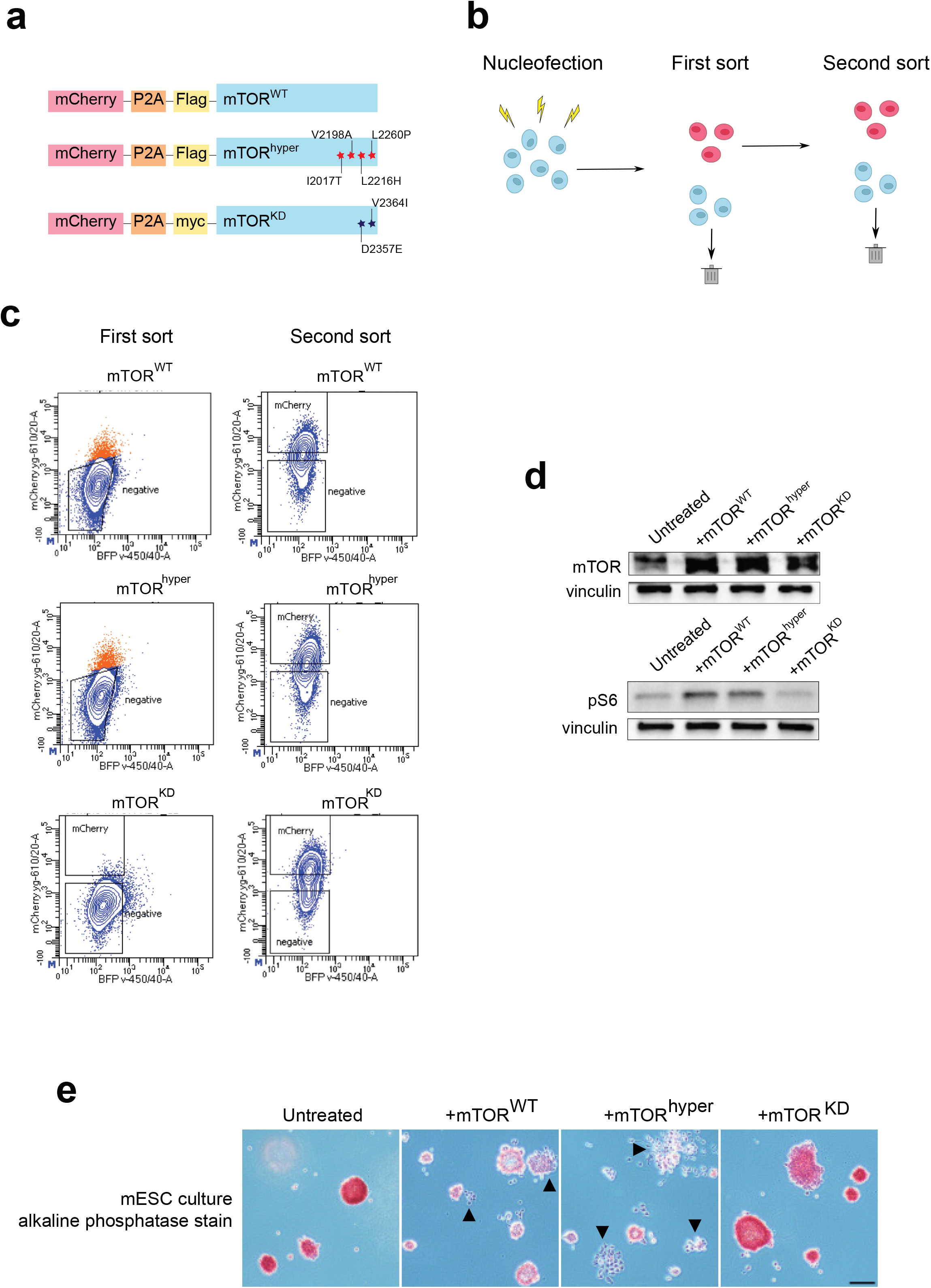
Ectopic overexpression of mTOR leads to increased differentiation of ESC colonies in serum/LIF culture conditions. a. Overexpression constructs for wild-type, hyperactive, catalytic-dead mTOR. The corresponding mutations are indicated. b. Experimental workflow. Cells were cultured in serum/LIF-containing ESC media and sorted for mCherry expression twice to overcome silencing of the construct c. Flow cytometry plots from first and second sorts of mTOR^WT^, mTOR^Hyper^, and mTOR^CD^ cells. d. Western blots showing the expression levels of mTOR and phosphorylation of its downstream target ribosomal protein S6. Vinculin serves as loading control. e. Alkaline phosphatase staining of wild-type ESCs and ESCs overexpressing wild-type, hyperactive, or catalytic-dead mTOR. mTOR overexpression results in precocious differentiation.

**Figure S3.**
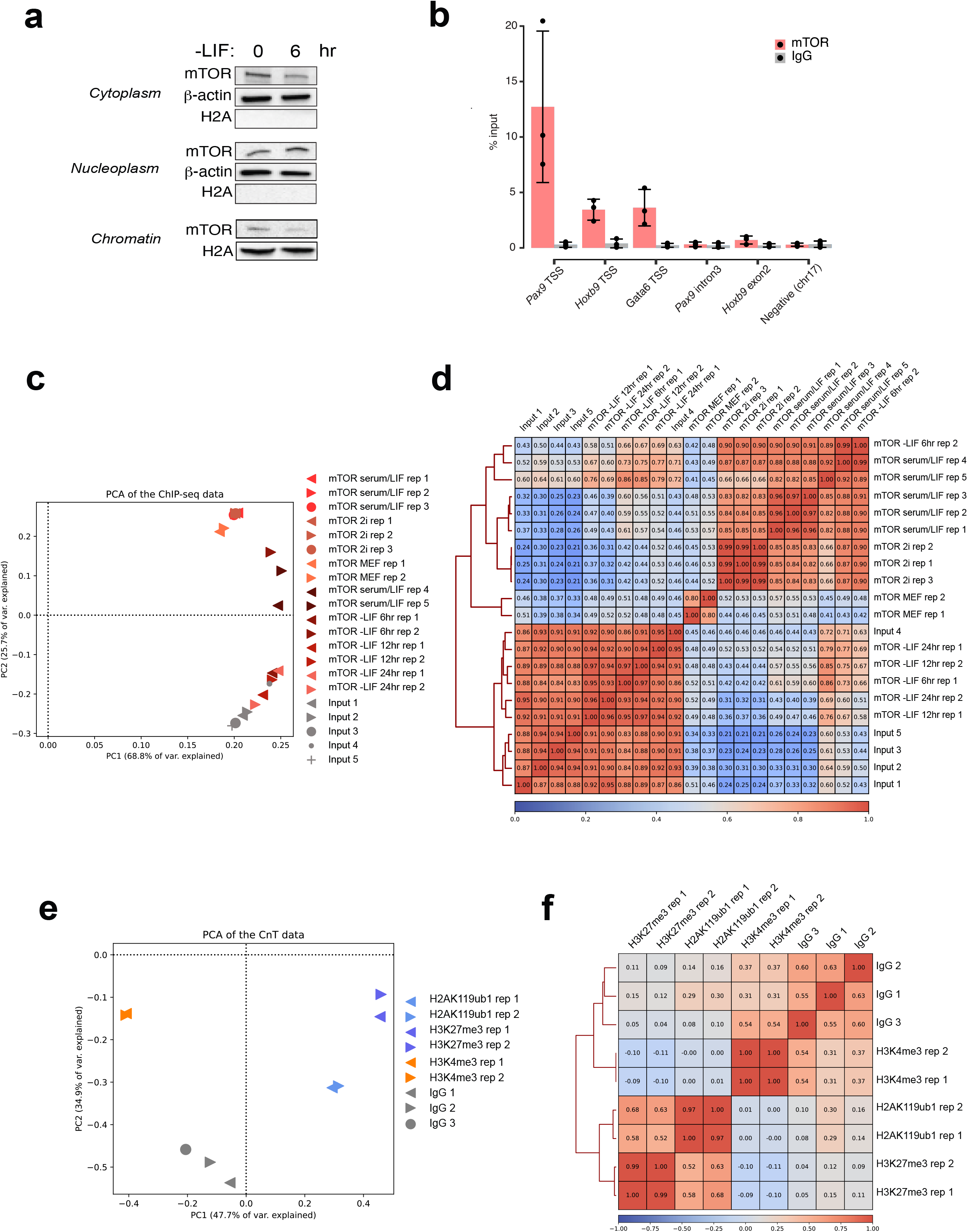
Quality assessment of ChIP-seq and CUT&Tag datasets. a. Western blot showing mTOR in different cellular fractions in ESCs cultured in serum/LIF or 6h after LIF withdrawal. b. ChIP-qPCR of mTOR binding at target genes along with a negative control region. c. Principal component analysis (PCA) based on the mean signal of the CPM normalized ChIP-seq samples at gene promoters (TSS +/-1kb). d. Correlation plot of all mTOR ChIP-seq and input samples based on the mean signal of the CPM normalized ChIP-seq samples at gene promoters (TSS +/-1kb). e. PCA based on the mean signal of the normalized CUT&Tag samples at gene promoters (TSS +/-1kb). f. Correlation plot of the shown CUT&Tag samples based on the mean signal of the CPM normalized ChIP-seq samples at gene promoters (TSS +/-1kb).

**Figure S4.**
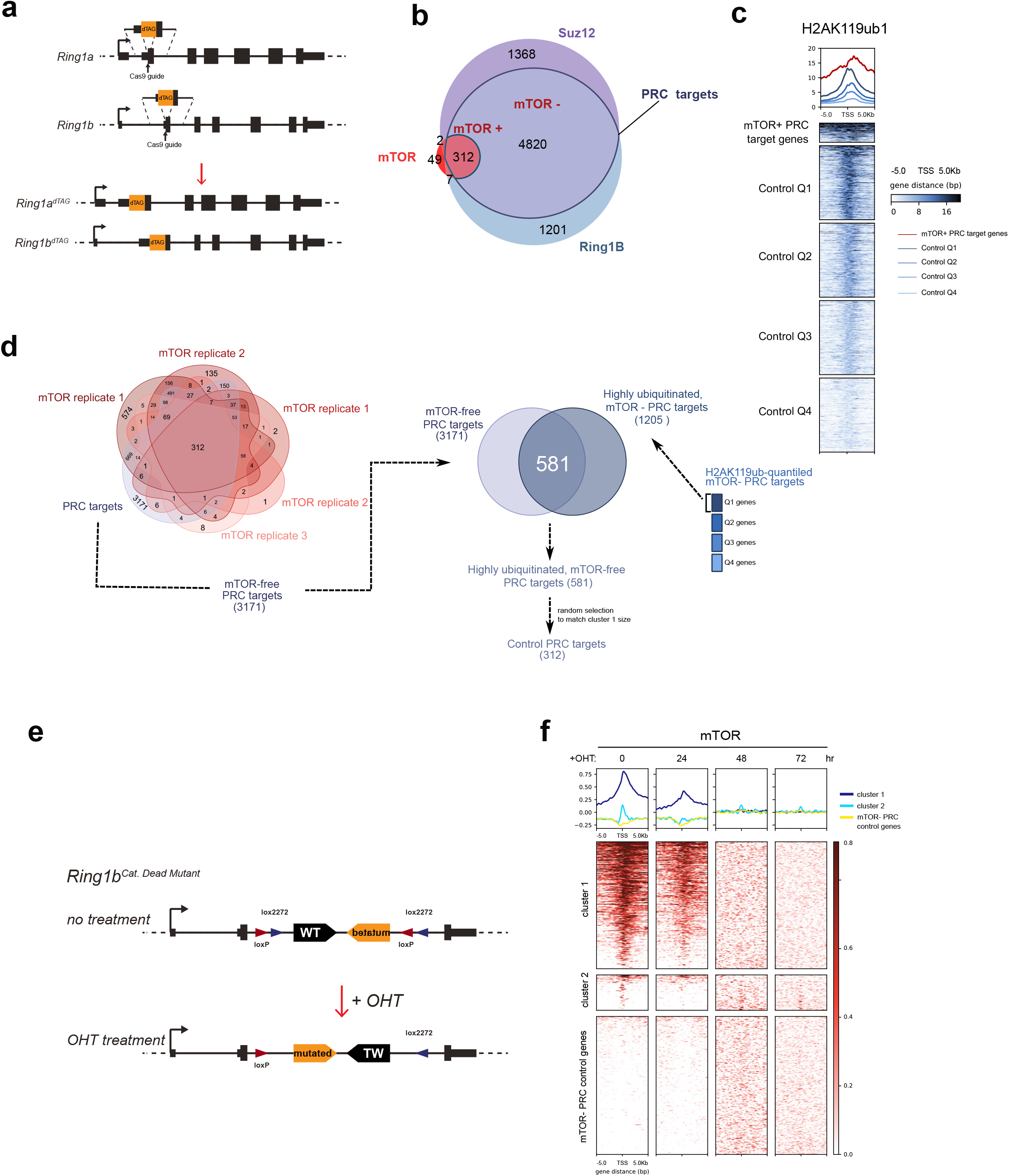
Further characterization of mTOR-PRC interaction. a. Schematic showing the strategy for generation of the *Ring1a/1b*^*dTAG*^ cell line. For both *Ring1a* and *Ring1b*, Cas9 guides specific for the 5’ end of each gene were used to insert the dTAG coding sequence via homology-directed repair. b. Venn diagram showing overlap of mTOR and PRC target genes (Suz12 and Ring1B targets). c. Heatmap showing H2AK119ub1 levels +/-5kb of TSS of mTOR target genes and the remaining PRC+ genes divided into four quantiles based on their H2AK119ub1 levels. d. Schematic showing the workflow of generating the mTOR-PRC+ control genes. e. Schematic showing the strategy for generation of the inducible PRC1 mutant cell line. f. Heatmap showing mTOR signal upon induction of PRC catalytic mutation +/-5kb of TSS of at mTOR target and mTOR-PRC+ control genes.

**Figure S5.**
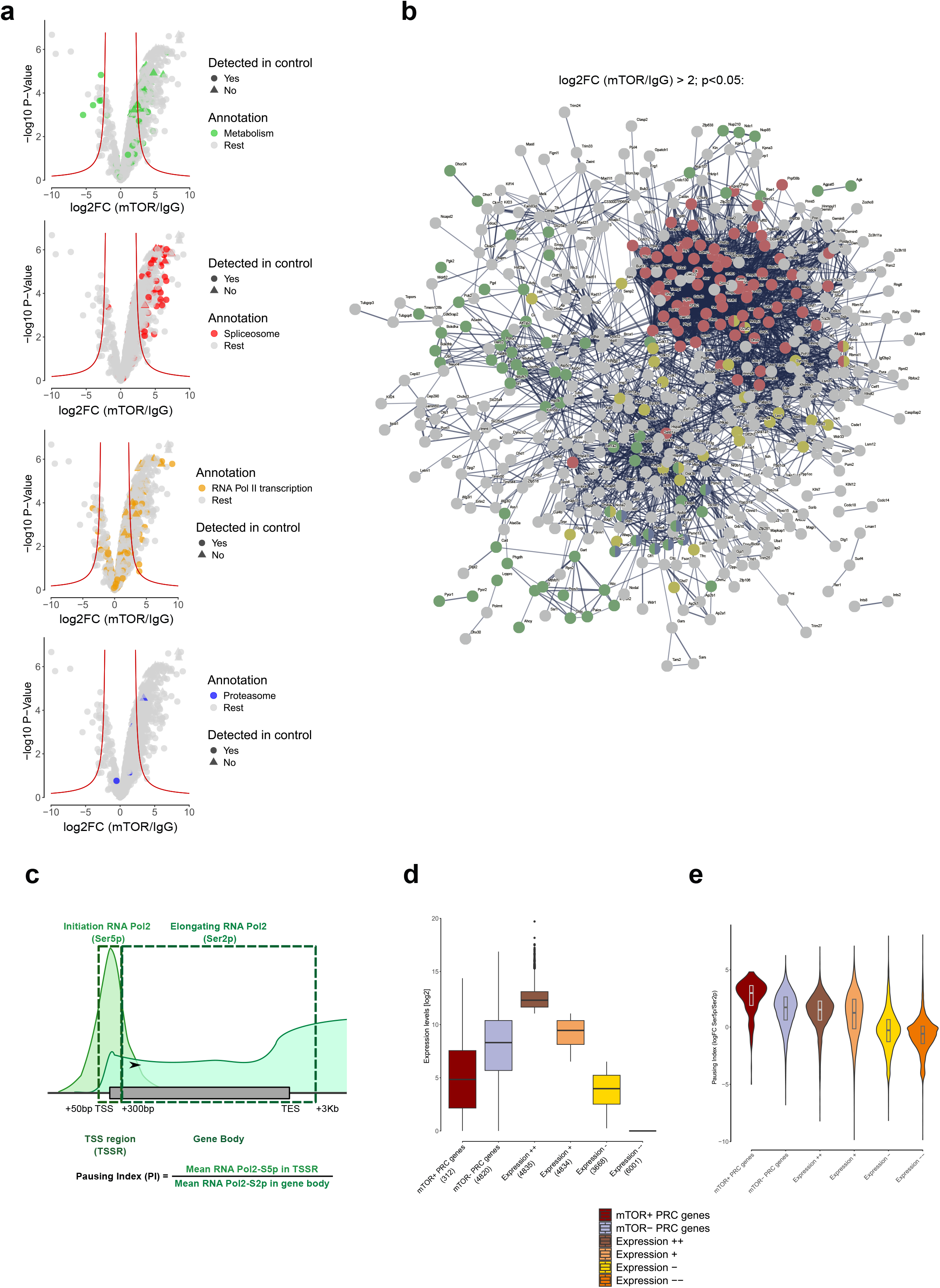
Further characterization of IP-MS and RNA Pol2 dynamics at mTOR target genes. a. Volcano plots the proteins belonging to each enriched pathway in mTOR IP-mass spec data. b. STRING interaction plot showing all significant proteins. Same pathways are highlighted as in Figure 5d. c. Schematic showing the definition of Pol2 pausing index. The figure and definition are based on Day et al^50^ but adjusted to consider the different states of the RNA-Pol2 instead of total levels. d. Box plot showing the nascent expression levels in untreated PRC1 degron ESCs of indicated gene sets. PRC-genes have been separated into expression quantiles. e. Violin plot showing pausing index distributions of the indicated gene groups.

